# Blood-Based Epigenetic Instability Linked to Human Aging and Disease

**DOI:** 10.1101/2025.02.25.640133

**Authors:** Salman Basrai, Ido Nofech-Mozes, Rajesh Detroja, Fernando L. Scolari, Mehran Bakhtiari, Andrea Arruda, Tracy Murphy, Scott V. Bratman, Steven M. Chan, Mark D. Minden, Jae-Sook Ahn, Dennis D. H. Kim, Robert Kridel, Filio Billia, Sagi Abelson

## Abstract

The abundance, dynamics, and context-dependent heterogeneity of DNA methylation—where a pattern considered abnormal in one cell type may be normal in another—poses challenges in identifying methylation abnormalities linked to disease risk. Through genome-wide analyses, we identified CpG sites with remarkably consistent methylation profiles in healthy whole blood, predominantly existing in an unmethylated state. We examined alterations at these epigenetically stable loci in diverse cohorts, including those with cardiovascular disease and hematological cancers. Our findings reveal methylation pattern disruption in myeloid and lymphoid malignancies, correlating with clonal burden fluctuations during leukemia treatment. In non-cancer cohorts, we observed that normally stable CpG sites exhibited progressive instability with advancing age, which was also associated with the onset of cardiovascular disease and decreased survival rates. This study links DNA methylation instability with the expansion of risk-prone blood cells and highlights its role as a biomarker for both cancer and cardiovascular disease.

## Main

Somatic cell evolution is driven by genetic alterations that enable cells to circumvent normal regulatory mechanisms governing proliferation and tissue organizations^1,2^. For these alterations to enhance clonal fitness, they must persist long enough to evade cell death and be heritable, thereby increasing the population of affected cells and facilitating the accumulation of additional drivers of clonal expansion. In extreme cases, the accumulation of genetic alterations can result in more extensive genomic instability and cancer. Consistent with these established principles, genetic mutations follow a paradigm in which they persist throughout the lifespan of a cell lineage if not corrected before cellular replication^3,4^.

Conversely, epigenetic modifications present a paradox: while they can be heritable across cell divisions, they are also influenced by environmental factors and intrinsic cellular cues, making them more reversible and malleable than genetic mutations^5^. These characteristics complicate the definition of epigenetic normalcy, thereby challenging our ability to determine how epigenetic variability may drive disease initiation and progression.

Clonal hematopoiesis of indeterminate potential is an age-related condition marked by the dominance of one or more genetically distinct clones of blood cells^6,7^. This phenomenon, often initiated by leukemia-associated gene mutations in hematopoietic stem and progenitor cells, is associated with increased risks of hematologic neoplasm development^8–10^, adverse cardiovascular events^11,12^, and overall mortality risk^8,13^. While the genetic underpinnings of clonal hematopoiesis have been extensively investigated^14^, non-mutational mechanisms governing the emergence and expansion of hazardous blood cell populations remain incompletely understood.

Recent research reveals that up to 50% of clonal expansions in human blood may lack identifiable genetic drivers^8,15–17^, suggesting that additional factors, including epigenetic modifications, may play a crucial role. The observation that clonal hematopoiesis confers similar health risks regardless of the presence of known genetic drivers underscores the importance of elucidating novel mechanisms driving its emergence and expansion^8,15^.

This study introduces the concept of DNA methylation instability, a phenomenon characterized by aberrant changes in methylation levels at typically unperturbed CpG sites. We identified epigenetically stable loci across the human blood methylome, explored the molecular underpinnings of their stability, and investigated the implications of their disruption in various conditions previously linked to clonal hematopoiesis.

## Results

### Identification of epigenetically stable CpG sites

To identify stable features of the human methylome, we analyzed whole blood DNA methylation profiles from a large cohort of young, healthy individuals where substantial clonal populations are expected to be infrequent (n=1,658, age=18)^18^. Genome-wide DNA methylation levels were assessed using Illumina 450K Infinium arrays. Methylation levels were calculated using the ratio of intensities between the methylated and the combined locus intensity (β-value). CpG sites were restricted to autosomes, excluding sites near common single nucleotide polymorphisms, those associated with probes known for ambiguous mapping or cross-reactivity^19^, and sites predictive of biological age^20,21^. Additionally, samples corresponding to abnormal blood cell count estimates, as determined by deconvolution analysis^22^, were excluded (Supplementary Fig. 1). This quality control process resulted in a final Discovery Cohort of 1,525 individuals.

We defined Epigenetically Stable Loci (ESLs) as the 10% least variable CpG sites based on methylation level variance across the cohort. Given that peripheral blood contains a mixture of cell types, consistent methylation levels among different cell populations were expected by this approach. We confirmed low variability in purified blood cell types^23^ and further refined our selection of ESLs by excluding CpG sites with outlier β-values in any of the cell categories. (Supplementary Fig. 2). This approach identified 31,569 unmethylated and 6,099 methylated ESLs (Fig. 1a,b).

**Fig. 1:**
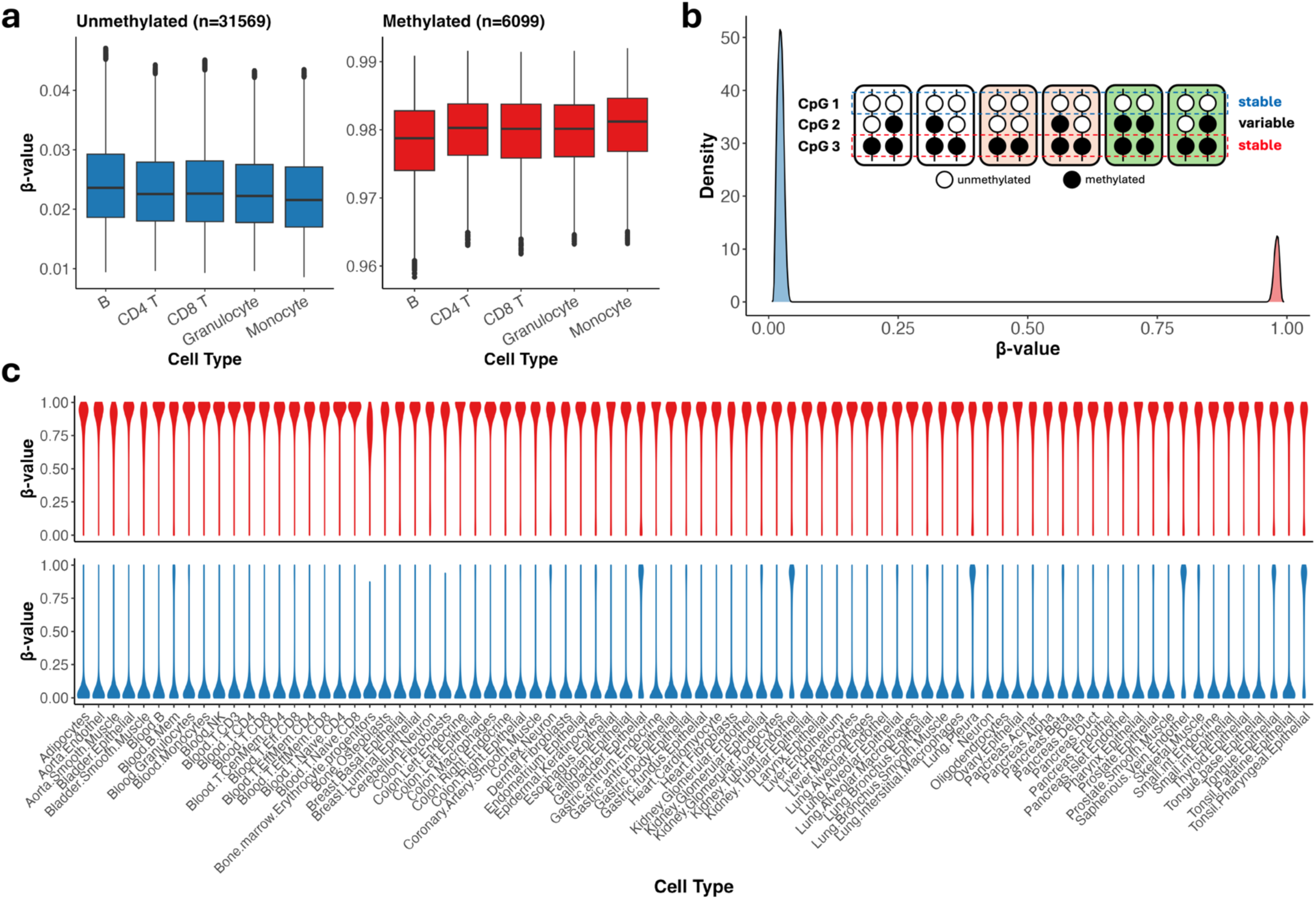
Epigenetically stable loci (ESLs) identified from peripheral blood of young, healthy individuals. **a**, Average β-values of unmethylated (left) and methylated (right) ESLs in purified blood cell populations (n = 28 samples per cell type). **b,** Density plot of average β-values for all ESLs across the Discovery Cohort. Blue shading indicates unmethylated ESLs, while red shading represents methylated ESLs. The diagram depicts an example of 6 diploid cells from 3 individuals. CpG 1 is stably unmethylated, CpG 2 is variable across individuals, and CpG 3 is stably methylated. **c,** Violin plots of β-values at methylated (top) and unmethylated (bottom) ESLs in purified populations of 82 human cell types from adult healthy tissues, derived from whole-genome bisulfite sequencing. For this dataset, β-values for each locus were calculated as the ratio of methylated reads to total reads.

Notably, although these ESLs were identified in blood, we also observed their characteristic extreme methylation levels—either highly methylated or unmethylated—in purified cells from non-hematological tissues^24^ (Fig. 1c). This consistency across diverse tissue types and detection techniques suggests that these extreme methylation levels at ESLs are not attributable to data analytic or technical artifacts associated with the Infinium arrays, but rather a reflection of stringent regulatory control at these sites.

### Epigenetic perturbation at normally stable loci is common in blood malignancies

Dysregulation of DNA methylation is a well-established hallmark of cancer^25,26^. Therefore, we hypothesized that the ESLs identified in healthy blood samples may be disrupted during carcinogenesis. We analyzed DNA methylation levels across a diverse spectrum of hematological malignancies and additional non-cancer controls (Supplementary Table 1). Our analysis revealed that, with the exception of chronic phase CML, all investigated hematological cancers displayed elevated methylation levels across the normally unmethylated ESLs. This pattern was evident across diverse cancers, irrespective of their classification as pediatric or adult cancers, biopsy tissue origin (i.e., bone marrow, blood, or lymphoid tissue), or the patient’s biological sex (Fig. 2).

**Fig. 2:**
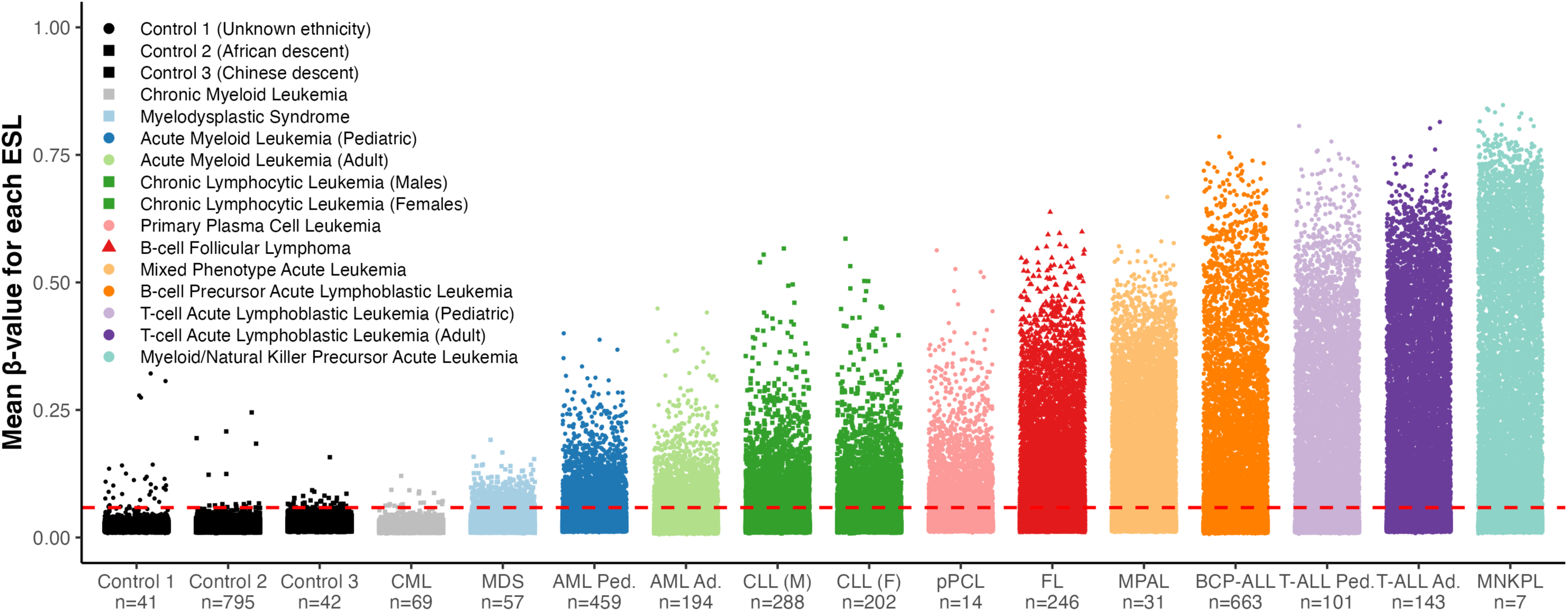
Destabilization of unmethylated ESLs in hematological malignancies. β-values of unmethylated ESLs across various cancer and control cohorts. For each cohort, the average β-value for each of the 31,569 ESLs is plotted, with each point representing a single ESL. The dotted red line indicates the destabilization threshold (β = 0.0586) determined from the control cohorts. The tissue source for each cohort is indicated by marker shape: circles for bone marrow, squares for peripheral blood, and triangles for lymphoid tissue. CML: Chronic Myeloid Leukemia, MDS: Myelodysplastic Syndromes, AML: Acute Myeloid Leukemia, CLL: Chronic Lymphocytic Leukemia, pPCL: Primary Plasma Cell Leukemia, FL: Follicular Lymphoma, MPAL: Mixed Phenotype Acute Leukemia, BCP-ALL: B-cell Precursor Acute Lymphoblastic Leukemia, T-ALL: T-cell Acute Lymphoblastic Leukemia, MNKPL: Myeloid/Natural Killer Cell Precursor Acute Leukemia, Ped: Pediatric, Ad: adult, M: male, F: female.

To assess inter-cohort differences, we established a destabilization threshold based on ESL β-values derived from the three independent control cohorts in this dataset (n = 878). This threshold was determined by calculating the mean β-value for each ESL in each control cohort, then compiling these means into a single list. The 99.9th percentile of this compiled list was set as the destabilization threshold (β-value = 0.0586).

Lymphoid cancers exhibit a significantly greater degree of epigenetic destabilization than myeloid cancers, reflected in both the number of perturbed ESLs as well as methylation levels at those loci. For example, the median number of perturbed ESLs in AML patients was 475 as compared with 4763 in T-ALL (Supplementary Fig. 3; *P* value < 0.001). The average β-value of perturbed ESLs in AML was 0.2934 as compared with 0.4623 in T-ALL (*P* value < 0.001).

We also observed hypomethylation of normally methylated ESLs. However, this pattern was less consistent across different cancers and exhibited greater variability among controls (Extended Data Fig. 1). Therefore, we focused on the 31,569 unmethylated ESLs in this study – any subsequent mention of ESLs refers exclusively to this subset.

### Association between chromatin status in precursor cells and ESL perturbation susceptibility

Given the observed differential degrees of epigenetic instability between myeloid and lymphoid cancers, we investigated the chromatin accessibility of ESLs in corresponding cells. To identify lineage-enriched perturbed ESLs, we compared AML patients’ profiles (n = 997) to T-ALL (n = 353) and BCP-ALL; n = 663) (Supplementary Table 1). We binarized the β-value matrix using the 99.9th percentile of non-cancer cohorts as a threshold (Supplementary Table 2), then performed an enrichment analysis.

This analysis revealed 1,072 ESLs that were predominantly perturbed in T-ALL and BCP-ALL, which were classified as lymphoid-enriched (*P* value < 0.001). In contrast, only 81 ESLs were identified as myeloid-enriched (*P* value < 0.001) (Supplementary Fig. 4). This disparity is in accordance with the overall lower degree of ESL hypermethylation observed in AML compared to lymphoid cancers (Fig. 2).

To explore the relationship between instability in these lineage-enriched ESLs and chromatin accessibility, we analyzed single-cell ATAC-seq profiles of healthy hematopoietic cells^27^. We computed chromVAR deviation scores for each cell (n = 22,737), separately for lymphoid-enriched and myeloid-enriched perturbed ESLs. These deviation scores quantify the extent of chromatin accessibility at specific regions, comparing observed accessibility to the dataset-wide expectation^28^.

Chromatin regions corresponding to lymphoid-enriched instability exhibited low accessibility in common lymphoid progenitors, T-cell and B-cell lineages, and relatively higher accessibility in hematopoietic stem cells (HSCs), common myeloid progenitors, and granulocyte-monocyte progenitors. Conversely, myeloid-enriched perturbed ESLs exhibited significantly reduced chromatin accessibility in HSCs and myeloid progenitors, but relatively increased accessibility in lymphoid-lineage-associated cells. (Fig. 3; *P* value < 0.001). These contrasting patterns link lineage-associated ESLs and their frequent perturbation in leukemia to differential chromatin accessibility in corresponding healthy cell types. We further confirmed these associations in a second dataset^29^ (Extended Data Fig. 2).

**Fig. 3:**
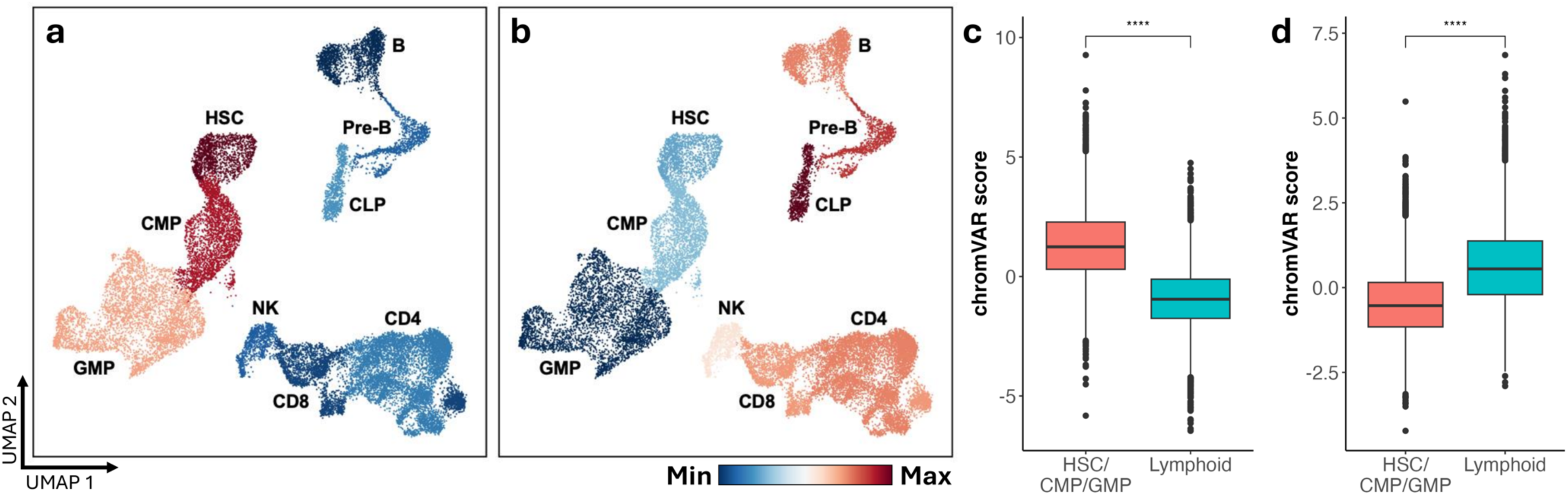
Chromatin accessibility patterns in normal cells associated with lineage-enriched ESLs. **a**, **b**, Chromatin accessibility UMAPs of 22,737 hematopoietic cells from healthy donors. Each cell is colored according to the average chromVAR deviation score for its respective cell type, based on chromatin accessibility at specific genomic regions corresponding to (**a**) lymphoid-enriched ESLs and (**b**) myeloid-enriched ESLs. Red shading indicates less compacted chromatin (higher accessibility), while blue shading indicates more compacted chromatin (lower accessibility). **c**, **d**, Comparison of HSCs, CMPs, and GMPs to lymphoid cell types (CLP, Pre-B, B, CD4 T, CD8 T, NK) based on chromVAR deviation scores corresponding to (**c**) lymphoid-enriched ESLs and (**d**) myeloid-enriched ESLs. **** p < 0.0001 for Wilcoxon test.

### Evidence of clonal epigenetic memory

We conducted an analysis of a pan-hematological cancer cohort comprising 3,019 samples (Supplementary Table 1) to characterize the frequency of destabilization events (Supplementary Table 2). Our findings revealed a significant positive correlation between the proportion of samples in which each ESL is destabilized and the average methylation level of the ESL in those destabilized samples, both across the entire pan-cancer cohort (r = 0.79, *P* < 0.001) and within individual cancer types (Supplementary Fig. 5). Specifically, ESLs destabilized in a higher proportion of samples showed higher average methylation compared to those destabilized in fewer samples. This pattern suggests that when an ESL is destabilized more frequently across patients, it often involves a larger proportion of cells within each patient as well.

The recurrent perturbation of identical ESLs across patients, coupled with the well-established epigenetic plasticity of DNA methylation^5^, suggests one explanation for the elevated signals observed in cancer: different cells within a single patient may independently acquire methylation in the same ESL. An alternative explanation and contributing factor is the heritability of the methylation signal. To investigate this, we sought evidence of somatic methylation inheritance at ESLs. Specifically, we defined low-recurrence ESLs as those perturbed in fewer than 5% of patients in the pan-hematological cancer cohort (Supplementary Table 2). This approach helped identify ESLs that are more likely to reflect unique epigenetic alterations specific to individual patients.

Using datasets representing paired diagnosis and relapse leukemia samples (Supplementary Table 1), we identified 15 low-recurrence ESLs that exhibited the highest methylation levels in each patient’s relapse sample. We then performed unsupervised clustering to pair diagnosis samples with their corresponding relapse specimens from the same patients (Fig. 4a; Extended Data Fig. 3). In AML (n=15 diagnosis-relapse pairs), this approach resulted in 10 correct matches (Permutation test: *P* = 0.002). For chronic lymphocytic leukemia (CLL), 31 out of 40 pairs were correctly matched (*P* < 0.001). In BCP-ALL (n=24 pairs), all diagnosis samples were correctly assigned to their corresponding relapse sample (*P* < 0.001). The negligible probability of these results occurring by chance (also see Supplementary Note 1) provides compelling evidence for the clonal preservation of ESL methylation, suggesting epigenetic memory in leukemia cells throughout disease progression. This is further supported by the observed reduction in DNA methylation during remission, indicating that this signal is specific to leukemic clones (Fig. 4b, c). Our findings further indicate that therapy-resistant epigenetic clones (epi-clones) may pre-exist in patients prior to treatment initiation. Comparable pairing was observed when analyzing the 15 most destabilized low-recurrence ESLs identified in diagnostic samples (Supplementary Fig. 6).

**Fig. 4:**
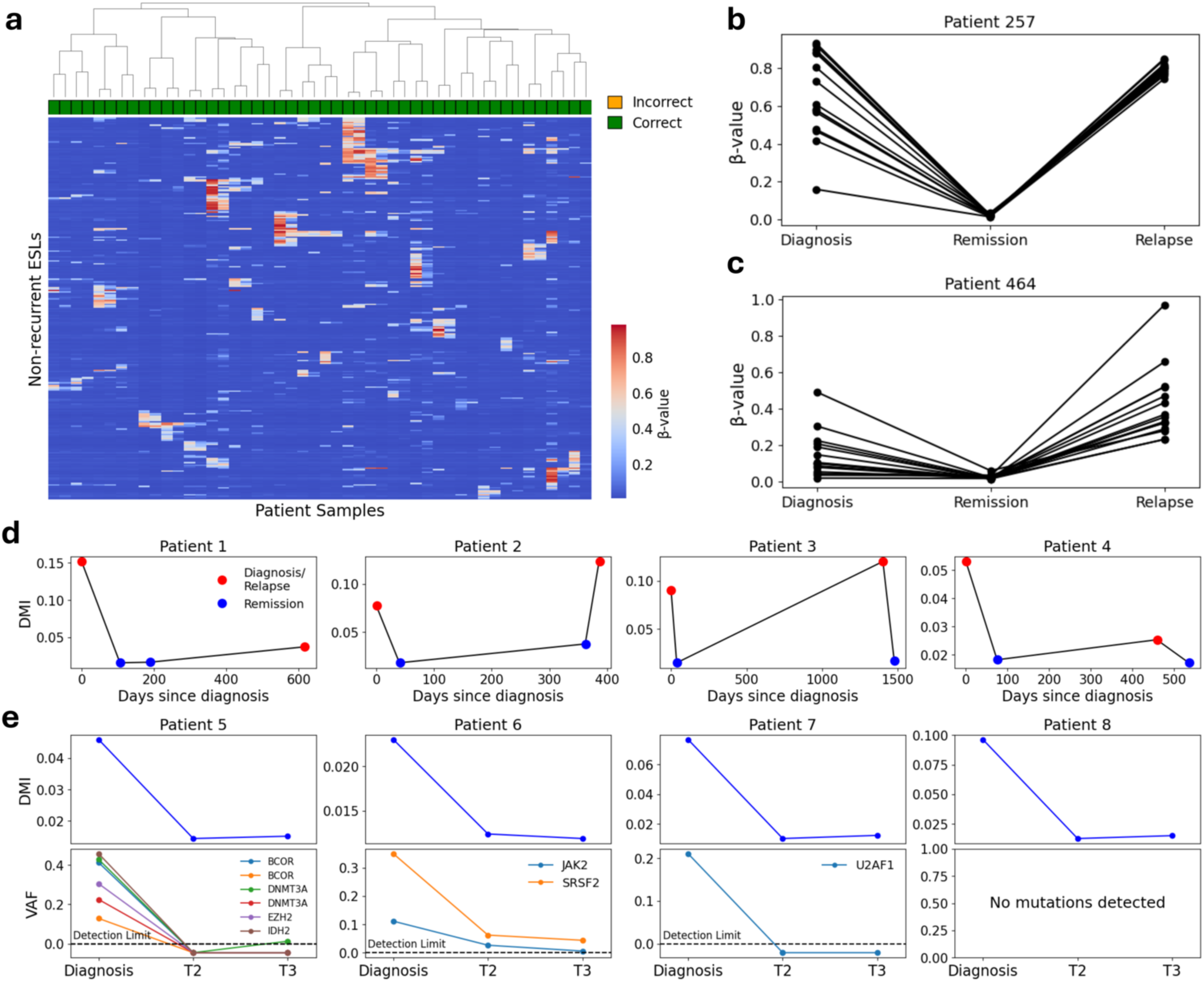
Aberrant DNA methylation in ESLs links leukemic epigenetic clones at diagnosis and relapse. **a**, Hierarchical clustering of paired diagnosis and relapse samples from 24 BCP-ALL patients. The top 15 most destabilized ESLs with low recurrence (i.e., perturbed in <5% of the pan-hematological cancer cohort) were selected from each patient’s relapse sample for clustering, resulting in a total of 273 unique ESLs examined across 48 samples. Correctly paired diagnosis-relapse samples are indicated in green. **b**,**c**, β-values of the top 15 low-recurrence destabilized ESLs in 2 BCP-ALL patients at diagnosis, remission, and relapse samples of which all 3 time points were available. Each line represents the β-value of a single ESL across the different disease stages. **d**, DNA methylation instability (DMI) levels in bone marrow samples from four AML patients (n = 4). Each patient was sampled at four distinct time points during the course of their disease. **e**, Analysis of peripheral blood samples from AML patients at diagnosis and two subsequent time points at remission. The top panels display DMI levels, while the bottom panels show variant allele frequencies (VAFs) of the patients’ leukemic mutations

### Temporal variability in DNA Methylation Instability

To quantify the degree of ESL perturbation on a per-sample basis, we introduced a metric called DNA methylation instability (DMI), defined as the standard deviation of β-values for ESLs within each sample. Since the majority of ESLs remain unperturbed within individual samples (Supplementary Fig. 3), we focused on ESLs with a perturbation rate above 5% across samples in the pan-hematological cancer cohort (n = 9,191; 29.1% of all ESLs, hereafter referred to as recurrently perturbed ESLs). This approach aims to increase the variability of DMI values across samples. Longitudinal measurements of bone marrow samples from AML patients (n = 4) revealed that DMI levels correspond to the expected patterns of leukemic burden across different clinical stages of the disease. Specifically, elevated DMI was observed at diagnosis and relapse, with lower levels during the first and second remissions (Fig. 4d). Notably, although Patient 2 was determined to be in remission at the third sampling, we observed an increase in DMI, potentially signaling the impending relapse.

In a previous study, we performed next-generation sequencing of AML patients at diagnosis and two subsequent time points post-therapy^30^. DNA methylation profiling of a subset of these patients (n = 10) revealed that DMI levels mirrored the allelic burden of previously identified somatic mutations. Specifically, DMI was elevated at diagnosis and decreased dramatically following induction therapy (Fig. 4e, Extended Data Fig. 4). These findings underscore the potential utility of monitoring DMI as a complementary approach to traditional mutation-based methods for assessing complete remission and residual disease post-treatment. This approach is particularly valuable in cases such as Patient 8, where clonal tracking was hindered by the absence of identifiable mutations.

### Increasing Instability with Advancing Age

Given that DMI measurements share similarities with genetic clonal markers in the hematopoietic system – particularly in their association with leukemic cell burden and the persistence of the alteration (Fig. 4, Extended Data Fig. 4) – and considering that hematopoietic clonality increases in frequency with age^31–33^, we hypothesized that DMI may also reflect age-related changes in individuals without hematological malignancies. To test the relationship between DMI and age, we interrogated several normal population cohorts of blood donors (Supplementary Table 1) and found significant correlations in all cases (Fig. 5a, Extended Data Fig. 5). These results demonstrate that the global deviation from the hypomethylated state of ESLs progressively increases with age, suggesting that DMI may be a general feature of aging hematopoietic cells rather than a phenomenon restricted to malignant transformation. Notably, we consistently observed lower DMI levels in these non-malignant cohorts (Fig. 5a and Extended Data Fig. 5) compared to leukemia diagnosis samples (Fig. 4d, e and Extended Data Fig. 4), indicating an accelerated rate of instability following malignant transformation.

**Fig. 5:**
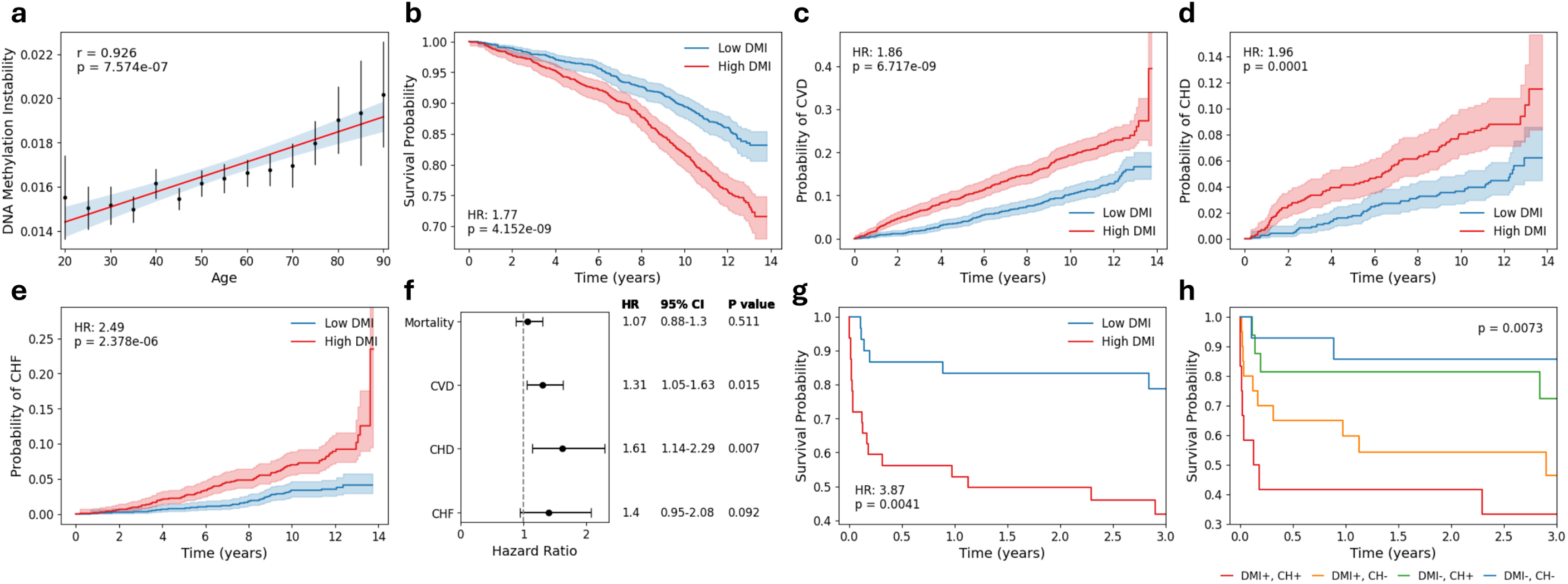
DNA Methylation Instability as a risk factor for cardiovascular disease, related mortality, and its association with age. **a,** DNA methylation instability (DMI) levels in a cohort of 656 healthy individuals. Participants were grouped into 5-year age categories. Black dots represent the mean DMI value for each age group, with error bars indicating the 95% confidence interval. Linear regression was performed on the group means. The blue shading represents the 95% confidence interval of the regression line. **b-e,** Univariate Kaplan-Meier analyses of 2,281 participants in the Framingham Heart Study (Exam 8) for **b**, mortality, **c**, cardiovascular disease (CVD), **d**, coronary heart disease (CHD), and **e**, congestive heart failure (CHF). Participants were classified as low DMI or high DMI based on the cohort median split. Shaded areas represent the 95% confidence interval. Hazard ratio (HR) and P values are indicated. **f,** Forest plot showing the effect of high DMI on each endpoint (**b-e**) using multivariate Cox proportional hazards regression, with correction for age and biological sex. **g,** Survival analysis of 64 cardiogenic shock patients, stratified into low DMI or high DMI groups based on a median split. **h,** Survival curves of the same patients, further stratified into the following categories: i) high DMI with presence of clonal hematopoiesis (CH) mutations, ii) high DMI with absence of CH mutations, iii) low DMI with presence of CH mutations, and iv) low DMI with absence of CH mutations. *P* values were calculated using the multivariate log-rank test.

### DNA methylation instability is associated with increased cardiovascular risk

Clonal hematopoiesis of indeterminate potential has consistently been linked to an elevated risk of cardiovascular diseases and related adverse outcomes, as demonstrated by our previous research and that of others^11,12,34–36^. Given the similar associations observed between the genetic definition of clonality and DMI in relation to both cancer and aging, we hypothesized that DMI could also serve as an indicator of cardiovascular risk.

To investigate this relationship, we utilized DNA methylation array data from the Framingham Heart Study Offspring cohort encompassing long-term follow-up data spanning up to 14 years^37^. Our objective was to determine whether DMI could stratify individuals into high-risk and low-risk groups for cardiovascular disease development. We employed Cox proportional hazards models to assess the relationship between DMI and several health outcomes, including all-cause mortality, incidence of cardiovascular disease (CVD), coronary heart disease (CHD), and congestive heart failure (CHF). Univariate survival analysis revealed that individuals with high DMI had a significantly higher risk for all of the defined endpoints compared to those with low DMI (Fig. 5b- e). After accounting for age and sex, the hazard ratios predicting elevated CVD and CHD risks remained significant (Fig. 5f).

In a previous study, we performed next-generation sequencing on patients with cardiogenic shock, a condition associated with early mortality. Our findings revealed significant differences in survival between patients with clonal hematopoiesis driven by mutations in epigenetic regulator genes, primarily *TET2*, and those without detectable clonal hematopoiesis^34^. To investigate the potential prognostic value of DMI in these high-risk individuals, we profiled a subset of patients from the original cohort, including 29 patients with mutations in either *DNMT3A* or *TET2* and 35 without clonal hematopoiesis. As a result, we deliberately obtained an underpowered cohort with respect to the association between clonal hematopoiesis and mortality rates seen in our original study (Supplementary Fig. 7). Nevertheless, individuals with high DMI demonstrated significantly lower survival compared to those with low DMI (Fig. 5g). This association remained significant after adjusting for age and clonal hematopoiesis status (HR: 3.76; *P* value = 0.006). Stratifying patients by both DMI levels and clonal hematopoiesis status revealed that DMI is an independent risk factor (Fig. 5h). This observation is further supported by our findings in AML diagnosis samples (Fig. 4e and Extended Data Fig. 4), where elevated DMI levels were consistently observed in patients with diverse genomic mutations, including those in the frequently mutated CH driver gene, *DNMT3A*.

### DNA methylation instability and gene regulation

The associations between DMI and aging (Fig. 5a, Extended Data Fig. 5), cancer (Fig. 2 and 4, Extended Data Fig. 3 and 4, Supplementary Fig. 3-6), and cardiovascular disease (Fig. 5b-h), prompted us to explore the underlying mechanisms that may drive these associations. Given the critical role of DNA methylation in gene expression regulation, we hypothesized that the gradual accumulation of epigenetic instability may disrupt gene expression and cellular function, potentially creating a permissive environment for the expansion of detrimental blood cells and the subsequent development of disease. Consistent with this hypothesis, we found that ESLs are significantly enriched within CpG islands compared to the other, non-ESL CpG sites targeted by the Infinium array (*P* value < 0.001; Fig. 6a), and are preferentially located in gene promoters, with 92.6% of ESLs occurring within 1500 bp of a transcription start site (TSS) (Fig. 6b).

**Fig. 6:**
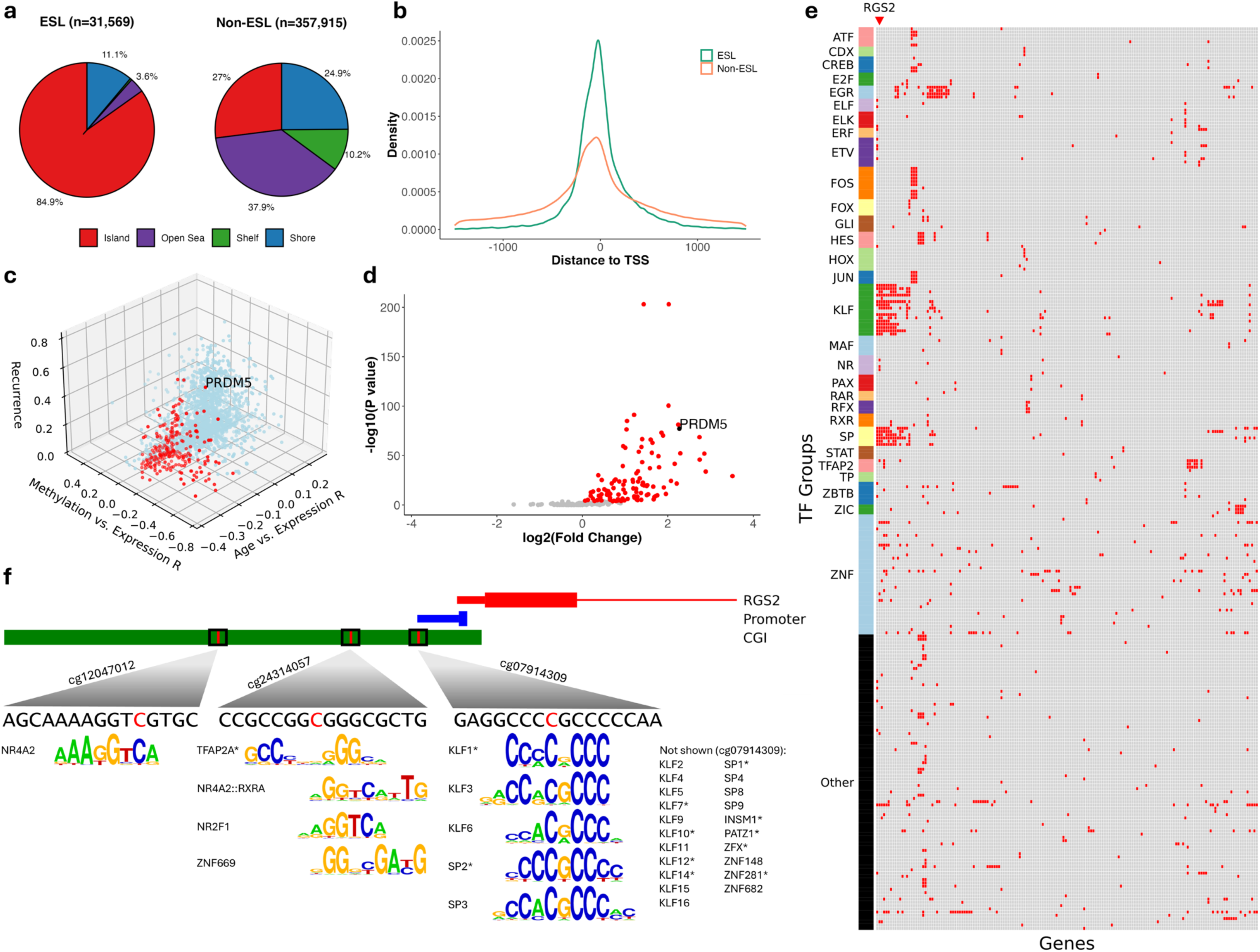
DNA methylation instability in promoter regions and its influence on gene expression. **a**, Genome distribution of ESLs (left) and non-ESLs (right). Each site is classified based on its proximity to CpG islands: within a CpG island, in a shore (0–2 kb from an island), in a shelf (2–4 kb from an island), or in the open sea (≥4 kb from an island). **b**, Density plot of distance to nearest transcriptional start site for all ESLs (green) and non-ESLs (orange). **c**, 3-Dimensional scatterplot of 1,879 genes – the intersection of three different analyses. The z-axis represents the mean recurrence of all ESLs within the promoter region of a given gene. The y-axis represents the mean Pearson’s correlation coefficient for ESL β-value vs. transcripts per million (TPM) of the corresponding gene in a pan-cancer dataset from The Cancer Genome Atlas (n = 8,960). The x-axis represents the Pearson’s correlation coefficient for TPM vs. age in a dataset of peripheral blood samples of healthy donors from the Genotype-Tissue Expression Project (n = 755). **d**, Volcano plot showing differential methylation analysis of promoter-associated CpG islands for 234 age-dependent genes. The y-axis represents the negative log-transformed p-values (-log₁₀(*P* value)), and the x-axis shows the log₂ fold change (log₂FC). Significant positive fold change values (Bonferroni-corrected *P* value < 0.05) are highlighted in red, indicating elevated methylation in AML compared to CD34+ control samples. **e,** Heatmap of age-dependent genes and transcription factor motifs in their promoter-associated CpG islands. Red boxes indicate intersections between the ESL and transcription factor binding motifs for the genes, highlighting regions where these elements overlap. **f**, Visualization of genomic features upstream of *RGS2*. The CpG island, promoter, and first exon are shown. For the promoter element, the “thin” section represents the 49 bp region upstream of the annotated transcription start site (TSS), while the “thick” section represents the TSS plus 10 bp downstream. Red lines on the CGI indicate ESLs. Transcription factor binding motifs containing the ESLs are shown.

Among the 1,879 genes with at least one recurrently perturbed ESL within their CpG island associated with the promoter, we identified 234 genes (12.5%) that are significantly downregulated in the blood of aging individuals^38^ (Fig. 6c, Supplementary Fig. 8). Given the complex interplay of multiple mechanisms regulating gene expression, we conducted a pan-cancer analysis using samples from The Cancer Genome Atlas (TCGA). This approach provided the necessary variability in both ESL methylation and gene expression levels to establish correlations between these variables, reinforcing the idea that perturbations in ESLs significantly influence the expression of these genes (Fig. 6c, Supplementary Table 3).

Based on ESL perturbation recurrence, the gene most frequently affected by DMI in that list was the tumor suppressor *PRDM5* (Supplementary Table 3). *PRDM5* has demonstrated growth-suppressive properties in experimental settings^39^ and is epigenetically silenced in various human cancers^39–41^. In this context, we observed a predominant trend of hypermethylation in AML for the 234 age-dependent genes, including *PRDM5 (*Fig. 6d).

Analysis of the gene list using g:Profiler^42^ revealed enrichment of transcription factor binding motifs, including those corresponding to SP and AP-2 families^43,44^, as well as other key regulators of cell cycle, differentiation, proliferation, and apoptosis (Supplementary Table 4). We confirmed that the CG bases of ESLs are integral to many of the enriched transcription factor binding motifs (Fig. 6e). This analysis highlighted associations between ESLs and key transcription factor binding motifs, including those associated with KLF proteins. For example, dysregulation of *KLF6* signaling was shown to play a significant role in both cancer and cardiovascular diseases^45^. Its ability to regulate apoptosis, inflammation, and cellular responses to stress makes it a critical player in maintaining cellular homeostasis. Another gene highlighted by our analysis is *RGS2*, whose promoter region contains ESLs that are integral to several different transcription factor motifs (Fig 6f). Down-regulation of *RGS2* in blood mononuclear cells is closely linked to cardiovascular disease through its effects on blood pressure regulation, G protein-coupled receptor signaling, and cardiac hypertrophy^46^.

Given that direct obstruction of transcription factor binding by cytosine methylation is a prevalent mechanism of gene repression mediated by DNA methylation^47^, these results imply a critical role of DMI in regulation of age-dependent genes, including known tumor suppressors.

## Discussion

Clonal hematopoiesis (CH) is a condition characterized by the clonal expansion of blood cells derived from a single hematopoietic stem or progenitor cell that has acquired somatic mutations^7^. This phenomenon is age-related and is associated with an increased risk of hematological malignancies, cardiovascular disease, and all-cause mortality^6^. Large-scale genomic studies have identified somatic mutations in leukemia-associated genes as key drivers of CH, underscoring the genetic basis of this aberrant clonal expansion^8,15–17,48^. Crucially, these studies have also demonstrated that CH frequently occurs in the absence of known driver mutations, sparking interest in alternative mechanisms that may drive the expansion of clonal blood cell populations.

This study underscores the role of epigenetic instability, its association with aging, and its direct connection to the health risks commonly associated with CH. Our analysis uncovered a set of CpG sites, termed ESLs, exhibiting remarkably consistent methylation patterns across healthy young adults and normal blood cell types. This low variability extended to cells from other healthy tissues, indicating that the methylation state at these loci is established early in development and typically maintained throughout cellular differentiation.

We discovered that disruptions in the hypomethylated state of ESLs in blood, coupled with increased methylation variability across these loci—a phenomenon we term DNA methylation instability (DMI)—are associated with detrimental health implications. This connection is supported by multiple observations in our study, highlighting the importance of preserving the hypomethylated state of ESLs for maintaining stable and healthy hematopoiesis.

Destabilization of ESLs was observed across a broad spectrum of hematological malignancies, aligning with the widespread epigenetic dysregulation characteristic of cancer cells ^26,49^. Elevated DMI was evident at leukemia diagnosis and significantly decreased post-induction therapy, paralleling changes in leukemic mutations. Elevated DMI was also evident in the absence of detectable mutations, underscoring its potential to augment genetic mutation testing for improved diagnostic accuracy and prognostic evaluation.

Moreover, we identified distinct epi-clonal patterns, characterized by specific ESL perturbations unique to each patient. These patterns persisted from diagnosis to relapse, indicating an epigenetic memory that withstands the selective pressures of cancer treatment. This persistence highlights the potential heritability of methylation at ESLs, which is necessary for exerting long-term biological impact. Over time, these heritable epigenetic alterations accumulate, potentially priming healthy cells for malignant transformation prior to the onset of overt leukemia. In support of this, we observed elevated DMI levels that positively correlated with age, indicating that the increasing dispersion of methylation patterns at ESLs is a progressive age-related phenomenon. While previous studies have identified both linear and non-linear associations between DNA methylation at specific CpG sites, age, and CH^50,51^, our approach differs in several key aspects, including the genomic localization of ESLs, their expected methylation levels in non-clonal samples, and their characteristic low variability in methylation across healthy individuals. These distinctions emphasize the tight regulatory control at ESLs and highlight the likelihood of their disruption playing causal roles in biological processes.

In the context of unhealthy aging, elevated levels of DMI were associated with increased cardiovascular risk. We observed a hazard ratio for developing coronary heart disease similar to that reported for clonal hematopoiesis of indeterminate potential (CHIP) in other epidemiological cohorts^11,52^. Notably, we found an association between DMI and cardiac patients’ survival that remained independent of common CHIP mutations in *TET2* and *DNMT3A*, suggesting that DMI contributes to cardiovascular risk through mechanisms distinct from those of established CHIP drivers.

Our findings demonstrate an association between DMI and genes whose expression decreases with age, including those involved in regulating cell fitness, cancer development and the onset of cardiac disease. Notably, we observed an enrichment of binding sites of essential transcription factor within these genes’ regulatory regions, including SP, KLF, FOS, JUN, EGR, and AP-2. ESL perturbations often coincided with these motifs, suggesting that methylation changes could impair transcription factor binding. Notably, many of these transcription factors are critical regulators of cell fate, orchestrating processes such as proliferation, apoptosis, and differentiation. Dysregulation of their binding affinity through ESL perturbations could have far-reaching effects on cellular function, potentially influencing hematopoietic stem cell maintenance, lineage commitment, and susceptibility to various diseases. Further research is needed to elucidate the genome-wide occupancy patterns of these transcription factors in both young and older individuals, as well as in leukemia patients and across different blood cell types. Such insights could advance our understanding of age-related disease etiologies and potentially lead to novel risk assessment and diagnosis strategies.

Our study has several limitations that should be acknowledged. By examining myeloid versus lymphoid enriched perturbed ESLs, we revealed that the chromatin state of precursor cells may influence the susceptibility of lineage-biased ESLs to perturbations in transformed cells. However, we cannot definitively conclude that chromatin compactness is the primary factor governing ESL stability and perturbation. The complexity of epigenetic regulation suggests that multiple factors likely contribute to DMI.

In addition, by examining low-recurrence perturbed ESLs, we were able to define patient-specific epi-clones, demonstrating that their ESL perturbation patterns are unique and maintained from diagnosis to relapse. This observation provides evidence for the heritability of DNA methylation at these sites, analogous to how somatic mutations are maintained in cancer cell populations. However, we cannot conclude that heritability is the primary driver of the overall elevated DMI observed in cancer. Future studies should focus on elucidating heritable epigenetic changes at ESLs versus recurrent de novo perturbations that occur independently in different cells. Understanding these dynamics could provide a valuable framework for distinguishing functionally relevant ESLs from background instability, akin to the distinction between driver mutations and passengers.

This study was conducted by leveraging extensive datasets, primarily BeadChip array datasets, which interrogate approximately 450,000 CpG sites—representing less than 2% of the total CpG sites in the human genome—with a bias towards promoter regions. While the ESLs we identified are the most stable among the array-covered sites, they may not represent the most stable sites across the entire genome. This limitation highlights the potential of more comprehensive profiling to yield additional insights including ESLs and their perturbation in distal regulatory regions.

Beyond the links between DMI, human aging, and disease established in this study, our approach provides a conceptual framework that connects epigenetic stability to normalcy. This framework could be extended to investigate changes beyond DNA methylation, including other heritable yet dynamic epigenetic marks, such as histone modifications. Applying this concept to diverse age-related diseases linked to blood clonality has the potential to enhance diagnosis, risk stratification, and targeted interventions aimed at mitigating the impact of both clonal and epi-clonal hematopoiesis on human health.

## Methods

### DNA methylation profiling

Venous blood and bone marrow biopsies were collected from 14 AML patients, resulting in a total of 46 longitudinal samples. For patients experiencing cardiogenic shock (n=64), buffy coat samples were utilized for DNA methylation profiling. Genomic DNA was extracted using the QIAamp DNA Blood Mini Kit (Qiagen) according to the manufacturer’s instructions. The concentration and purity of the DNA were assessed spectrophotometrically on a Qubit Fluorometer, with criteria set at an A260:A280 ratio greater than 1.7 and an A260:A230 ratio greater than 1.7. DNA integrity was evaluated through 1% agarose gel electrophoresis, loading 50- 100 ng of DNA per sample. Samples exhibiting a clear single band above the 10 kb DNA ladder were deemed suitable for further processing. High-quality genomic DNA (500 ng) was bisulfite-converted using the EZ DNA Methylation Kit (Zymo Research) following the manufacturer’s protocol. DNA methylation profiling was conducted using the Infinium MethylationEPIC microarray (Illumina). Patients Samples were collected in accordance with procedures approved by the Research Ethics Board of the University Health Network (REB #01-0573) and viably frozen in the Princess Margaret Leukemia Tissue Bank. Blood biospecimens from cardiogenic shock patients were collected under UHN REB approval (#18-6188) from the Peter Munk Cardiac Centre Cardiovascular Biobank. Written informed consent was obtained from all patients in accordance with the Declaration of Helsinki. University of Toronto REB approval was obtained for the use of datasets in secondary data analysis under Protocol #00041924.

### Methylation array data preprocessing

We utilized the R Bioconductor package minfi^22^. Data preprocessing commenced with raw IDAT files, except for the original dataset by Hannum et al.^20^, where preprocessed methylation intensities were available. Each individual IDAT file was loaded into an RGChannelSet object using the ‘*read.metharray.exp*’ function. To ensure consistency across different array types, we employed the ‘*convertArray’* function to restrict our analyses to probes common to both the MethylationEPIC and HumanMethylation450k arrays. Single-sample background and dye-bias correction were performed using the ‘*preprocessNoob’* function. The ‘*getBeta’* function was then applied to calculate the β-values, which range from 0 to 1 and represent the methylation level for each CpG site.

### Identification of epigenetically stable sites (ESLs)

Whole blood DNA from 1,658 young individuals^18^ (age = 18) was used to identify CpG sites exhibiting low variability in methylation levels. First, the function **‘***estimateCellCounts’* from the minfi package was employed to deconvolute blood cell proportions in each sample. Samples whose estimated proportions for any of CD4+ T cells, CD8+ T cells, B cells, natural killer cells, monocytes, or granulocytes were outliers (defined as values that fall outside 1.5 times the interquartile range from the first or third quartile.) were excluded from further analysis. Next, a series of filters were applied to the list of probes on the array. Only probes interrogated by both the Infinium HumanMethylation450 BeadChips and EPIC BeadChips were considered. Detection *P* values for these probes were evaluated across all the samples in the Discovery Cohort using the *detectionP* function. Probes for which less than 95% of the cohort had a detection *P* value below 1×10^-10^ were excluded. Additionally, we filtered out CpH probes and those targeting CpGs associated with single nucleotide polymorphisms, either within the CpG site itself or within the probe sequence. This was accomplished using the ‘*dropMethylationLoci’* and ‘*dropLociWithSnps’* functions. We also excluded CpG sites with prior evidence of potential cross-hybridization^19^. Lastly, we removed probes corresponding to CpG sites previously associated with age-related methylation changes^20,21^ as well as those targeting CpG sites located on sex chromosomes. The remaining CpG sites were ranked based on their β-value variance across the individuals in the Discovery Cohort. The 10% least variable sites were classified as either unmethylated or methylated, with a particular focus on unmethylated sites in this study. Classification was based on whether the average methylation level of each site was below or above 50%. To further reduce residual technical noise and minimize potential bias from specific cell types, the methylation level at those CpG sites were examined in purified blood cell populations. β-values were examined in B cells, CD4+ T cells, C8+ T cells, granulocytes and monocytes^23^. Any CpG site exhibiting an outlier β-value (defined as values that fall outside 1.5 times the interquartile range from the first or third quartile.) in any cell type was excluded from further analysis. After these steps, we retained a final set of 6,099 stably methylated loci and 31,569 stably unmethylated loci (ESLs).

### ESLs perturbation recurrence across blood cancers

To determine whether an ESL was destabilized in a given patient, we established a threshold based on the ESL β-values observed across three control cohorts independent of the Discovery Cohort (Supplementary Table 1). For each control cohort, we calculated the mean β-value for each ESL individually, pooled all these values, and set the destabilization threshold at the 99.9th percentile Based on this approach, a threshold of β-values = 0.0586 was determined. ESLs with methylation levels surpassing this threshold were classified as perturbed, also referred to as destabilized. To assess the frequency of ESL destabilization, we compiled a pan-cancer dataset comprising 3,019 samples from various hematological malignancies, including AML, T-ALL, BCP-ALL, CLL, FL, DLBCL, pPCL, MNKPL, and MPAL. Notably, CML and MDS were excluded from this analysis due to their characteristic low burden of blast cells and ESL destabilization. A β-value matrix was generated, with CpGs as rows and patients as columns. This matrix was then binarized by assigning a value of 1 to β-values exceeding the predetermined ESL destabilization threshold (β-values = 0.0586), and a value of 0 to β-values below the threshold (Supplementary Table 2). For each ESL, the proportion of patients exhibiting destabilization was calculated by dividing the sum of the binarized values for that row by the number of all samples (n = 3,019). Low-recurrence ESLs were defined as those perturbed in less than 5% of the samples while recurrently perturbed ESLs were defined as those above or equal to the 5% value. In the analysis examining the correlation between ESL perturbation recurrence and their average destabilized β-value (Supplementary Fig. 5), the average β-value for each ESL was determined only from patients exhibiting destabilization at this site.

### Identification of lymphoid- and myeloid-enriched perturbed ESLs

A comparison was conducted between AML samples and T-ALL and BCP-ALL to identify ESLs whose destabilization shows a stronger association with myeloid or lymphoid leukemias. The matrix of binarized β-values was used to conduct a Fisher’s test to assess the number of destabilized samples among myeloid cancer patients compared to lymphoid cancer patients. We only retained ESLs with a *P* value < 0.05 following Bonferroni correction. To identify lymphoid-enriched perturbed ESLs, we first selected sites that had an odds ratio greater than 1 and exhibited destabilization in at least 50% of the combined T-ALL and BCP-ALL samples, thereby emphasizing ESLs that are frequent in these lymphoid leukemias. This set of ESLs was ranked by their odds ratios and the top 50% of these sites were designated as lymphoid-enriched ESLs – those most strongly associated with lymphoid cancers. To identify myeloid-enriched perturbed ESLs, we employed a slightly different approach to address the relatively lower frequency of ESL perturbations characteristic of AML. Comparisons between AML and T-ALL, as well as AML and BCP-ALL, were conducted separately and the intersection of both the resulting lists was designated as myeloid-enriched. Additionally, we retained only ESLs that exhibited destabilization in at least 10% of the AML samples.

### Chromatin accessibility

A cell-by-peak matrix was obtained from Granja *et al.*^27^, corresponding to peaks called from the aligned scATAC-seq fragments for each cell. In this original study, latent semantic indexing (LSI) followed by singular value decomposition (SVD) were used to perform dimensionality reduction, normalization, and selection of optimal features (see original publication for details). To interrogate the relationship between lineage enriched ESLs and chromatin accessibility, a subset of the LSI-SVD matrix was created including the following cell types: HSC, CMP, GMP, CLP, Pre-B cells, B cells, CD4 T cells, CD8 T cells, NK cells. UMAP was performed on this matrix using the implementation in the uwot R package, with the same parameters used in the original publication. In order to assess chromatin accessibility at the lymphoid-enriched and myeloid-enriched ESLs, we first mapped ESLs to peaks in the scATAC-seq dataset. For each ESL, a 200 bp window was constructed with the ESL’s CpG bases at the center. Any scATAC-seq peaks that overlapped with these windows were retained. This process was performed separately for lymphoid-enriched ESLs (n=1072) and myeloid-enriched ESLs (n=81), yielding two peak sets corresponding to these loci. ChromVAR deviation scores^28^ were generated for each peak set using the ‘*AddChromatinModule’* function from the Signac R package. Each cell type in the UMAP was colored according to its average chromVAR deviation scores for lymphoid-enriched ESLs and myeloid-enriched ESLs. The same approach was used for the analysis of a second single cell ATAC-seq dataset from Izzo et al^29^. In this dataset of hematopoietic cells derived from patients with JAK2^V617F^-mutated myelofibrosis, only cells that were wild type for this allele were analyzed (n = 6,088). UMAP coordinates provided in the original publications were used for visualization.

### Survival and time-to-event analyses

DMI was defined as the standard deviation of ESLs within a sample, as outlined in the main text, and was calculated for each participant in the Framingham Heart Study^37^ (Exam 8; n=2,724). Patients were stratified into two groups, “high DMI” and “low DMI,” based on the entire cohort’s median DMI value. This binary predictor variable was then used to perform Kaplan-Meier survival analyses using the *lifelines* Python package. Cox proportional hazards (CoxPH) regression analysis was performed with DMI, age, and biological sex as covariates. Endpoints assessed included overall mortality, incidence of cardiovascular disease, coronary heart disease, and congestive heart failure. In all these analyses we only considered patients who had not already experienced any of the endpoints before their blood sample was taken (n=2,281). The 95% confidence intervals were calculated using the log hazard method (conf_type=“log-log”). The same approach was followed to conduct a univariate analysis in the cardiogenic shock sub-cohort to assess the association between DMI and survival. In the multivariate CoxPH regression analysis for these patients, DMI, age, and clonal hematopoiesis status were included as covariates. Clonal hematopoiesis status was treated as a binary variable to indicate the presence or absence of somatic mutations (specifically DNMT3A or TET2) in this sub-cohort.

### Power Calculation

Prior to DNA methylation profiling and the analysis assessing the impact of DMI status on the survival time of cardiogenic shock patients, we conducted a power analysis to determine the required sample size of a sub-cohort that would provide sufficient statistical power to detect a significant effect. This analysis was performed using the ‘*ssizeEpiCont’* function from the powerSurvEpi R package. Over 10,000 iterations, we randomly assigned a fixed number of patients from the entire cardiogenic shock cohort from our original study^34^ to high DMI status, while an equal number were assigned to low DMI status, mimicking the process of splitting patients based on median DMI values. The analysis was designed to achieve 85% power, assuming a hazard ratio of 1.5 and a type I error rate (alpha) of 0.05. This simulation indicated that a sub-cohort of 64 patients would be adequately powered to examine the relationship between DMI status and survival time in these high-risk patients.

### Genomic Distribution of ESLs

The association of ESLs with CpG islands, shores, shelves, and open sea regions was extracted directly from the Illumina 450K bead-array manifest. Enrichment analysis for ESLs within these regional categories was performed using Fisher’s exact test. Additionally, genomic coordinates of TSS were obtained from the UCSC Genome Browser annotation track database (https://genome.ucsc.edu/cgi-bin/hgTables; specifically, “NCBI RefSeq” track from the “Genes and Gene Predictions” group). Promoter regions were defined as the area from 1500 bp upstream to 1500 bp downstream of a TSS. The enrichment of epigenetically stable loci within these regions was assessed using Fisher’s exact test, with non-stable CpG sites (n = 357,915) serving as the reference set.

### Gene expression analysis

To identify genes for which ESL perturbation is a significant factor in their expression, we conducted a pan-cancer analysis using harmonized DNA methylation and RNA-seq transcript per million (TPM) data from 8,960 solid cancer samples (TCGA Pan-Cancer dataset downloaded from https://xenabrowser.net/datapages/). We focused on ESLs located within 1,500 bp upstream of TSSs, resulting in 15,914 sites associated with 6,822 genes after intersecting with the TCGA data. For each ESL, Pearson’s correlation coefficient was calculated between the methylation level (β-value) and the expression (TPM) of the corresponding gene. For genes with multiple ESLs in their promoter regions, the mean correlation coefficient (r) was calculated. In a second analysis, the correlation between gene expression and age was calculated for each of the 6,822 genes using a dataset of healthy individuals from the Genotype-Tissue Expression (GTEx) project^38^ who had their peripheral blood assayed with RNA-seq. In a third analysis, we calculated the mean recurrence across ESLs within each gene promoter. These steps yielded three variables for each gene: methylation versus expression (r), age versus expression (r), and recurrence. A total of 234 genes were selected for further interrogation based on the following criteria: 1) a significant negative correlation between age and expression, 2) a significant negative correlation between methylation at promoter-related ESLs and expression, and 3) a mean ESL recurrence greater than 5%.

### Transcription factor binding motif analysis

Enrichment for transcription factor (TF) binding motifs was first investigated by using g:Profiler^42^ on the list of 234 age-dependent genes revealing enrichment of binding sites for specific TFs. In order to establish whether these TF’s binding sites overlapped with ESLs, the JASPAR Transcription Factor Binding Sites track from the UCSC Genome Browser was used. The list of all motif matches containing an ESL was filtered to only retain hits with a score above 400 (corresponding to *P* value < 0.0001), leaving 278 TFs with binding sites across 166 genes. A binary matrix was constructed, with transcription factors (TFs) as rows and affected genes as columns, to map which genes contained ESL-associated binding sites for each TF.

### Differential methylation analysis

CpG islands located in the promoter regions of the 234 age-dependent genes were analyzed using whole-genome bisulfite sequencing data from six genetically diverse AML samples (IDs: 412761, 868442, 400220, 545259, 548327, 573988) as reported in the original study^53^, along with six control samples (CD34+ cells from healthy donors). Differential methylation between the AML and control samples was assessed by comparing beta values for each CpG island and its associated ESLs using a Wilcoxon rank-sum test.

### Statistical analyses

Throughout the study, correlation tests were performed using Pearson’s correlation coefficient. For enrichment analyses, we employed Fisher’s exact test (e.g., Supplementary Fig. 4). The Wilcoxon rank-sum test was utilized to compare the number of perturbed ESLs and their methylation levels between AML and T-ALL, as well as for the comparisons of chromVAR deviation scores. All *P* values reported are two-sided. Where applicable, the Bonferroni correction for multiple hypothesis testing was applied. For the analysis pairing diagnosis and relapse samples based on low-recurrence, highly destabilized ESLs, a permutation test was performed to evaluate the probability of obtaining similar matching success by chance. For each cancer type dataset, 1,000 simulations were run in which 15 random low-recurrence ESLs were selected for each patient. The randomly selected collection of sites (n = 15 times the number of patients) was then used to perform unsupervised clustering, with Ward’s linkage method and Pearson’s correlation coefficient (r) as the distance metric. The *P*-value was calculated as the proportion of simulations in which the number of correctly paired patients was greater than or equal to the observed result, indicating the likelihood of obtaining such a result by chance.

## Data availability

The sources of publicly available DNA methylation datasets used in this study are listed in Supplementary Table 1. Whole genome bisulfite sequencing bigwig files for normal blood donors and AML samples were downloaded from https://wustl.box.com/v/wilsonIDHmethylation. Blood RNA sequencing from individuals without malignancies was sourced from the Genotype-Tissue Expression project, available at https://gtexportal.org/home/downloads (Analysis V8). Harmonized gene expression and DNA methylation data from The Cancer Genome Atlas (TCGA) was obtained from Xena available at https://xenabrowser.net/.

## Code availability

Data analysis was performed in Python (v. 3.11; https://www.python.org/) and in R (v. 4.2.2; https://www.r-project.org/). Accessory scripts developed during this study, including those used for generating the figures, are available at https://github.com/abelson-lab/DNA-Methylation-Instability.

## Acknowledgements

We sincerely acknowledge and thank all the patients who contributed biological samples, as well as the studies that made their data publicly available to support this research. We are also grateful to The Centre for Applied Genomics at The Hospital for Sick Children in Toronto, Canada, for their assistance with DNA methylation profiling. Our deepest thanks go to John E. Dick for his invaluable support in the cardiogenic shock and longitudinal AML studies, which provided critical datasets for this work, as well as for his review of the manuscript. We also thank Robert J. Vanner and Alex Murison for their thoughtful and critical review of the manuscript. This research was funded by an Investigator Award from the Ontario Institute for Cancer Research, supported by the province of Ontario (to SA). Additional funding was provided by the Canada Graduate Scholarship from the Canadian Institutes of Health Research and the Sona Naran Pancha Graduate Award in Leukemia Research (to SB).

## Author contributions

SB performed all data analyses, interpreted results, contributed to concept development, and co-wrote the manuscript. INM contributed to single-cell data analysis and sample processing. RD, FLS, MB, AA, JA and SVB facilitated sample and data acquisition and clinical data curation. TM, SMC, MDM, DDHK, RK, and FB managed patient recruitment, data and sample acquisition and provided clinical expertise. SA conceptualized and designed the study, contributed to data analysis, led and supervised all aspects of the study, and co-wrote the manuscript. All authors read and approved the manuscript.

## EXTENDED DATA FIGURES

**Extended Data Fig. 1:**
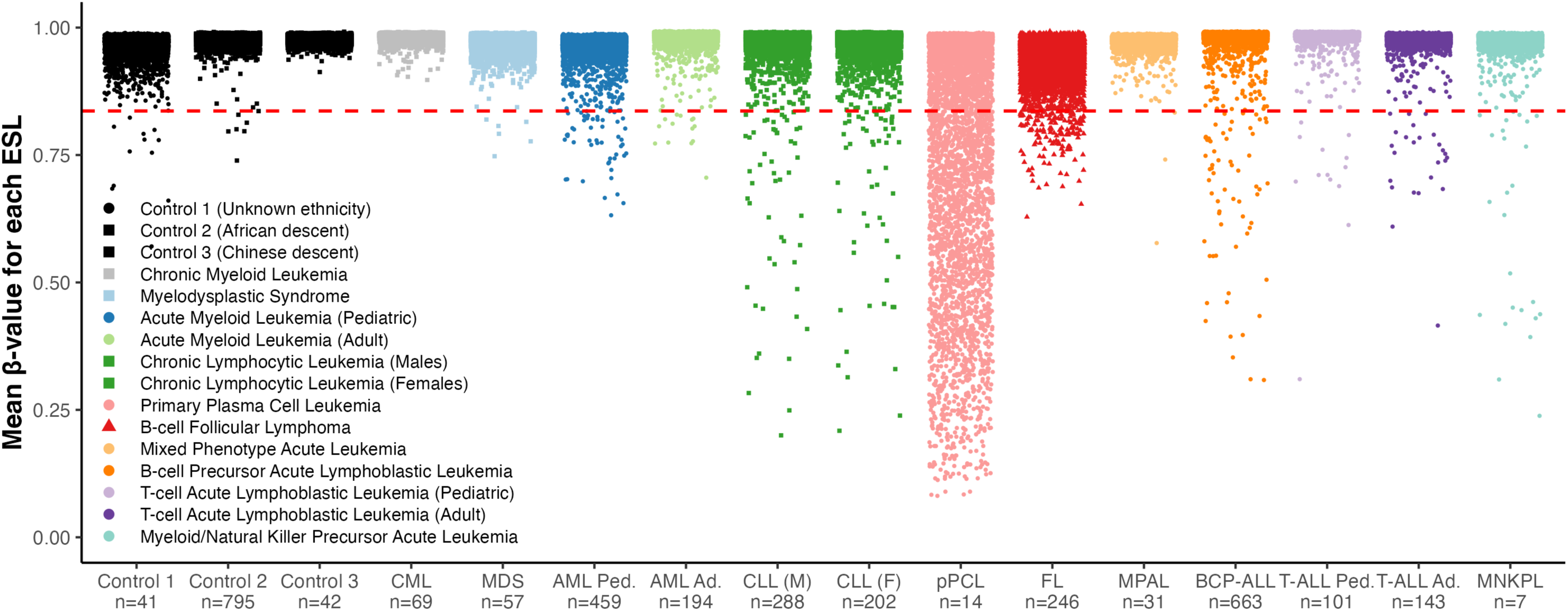
Destabilization of methylated ESLs in hematological malignancies. β-values of methylated ESLs across various cancer and control cohorts. For each cohort, the average β-value for each of the 6,099 sites is plotted, with each point representing a single ESL. The dotted red line indicates the destabilization threshold (β = 0.8364) determined from the control cohorts. The tissue source for each cohort is indicated by marker shape: circles for bone marrow, squares for peripheral blood, and triangles for lymphoid tissue.

**Extended Data Fig. 2:**
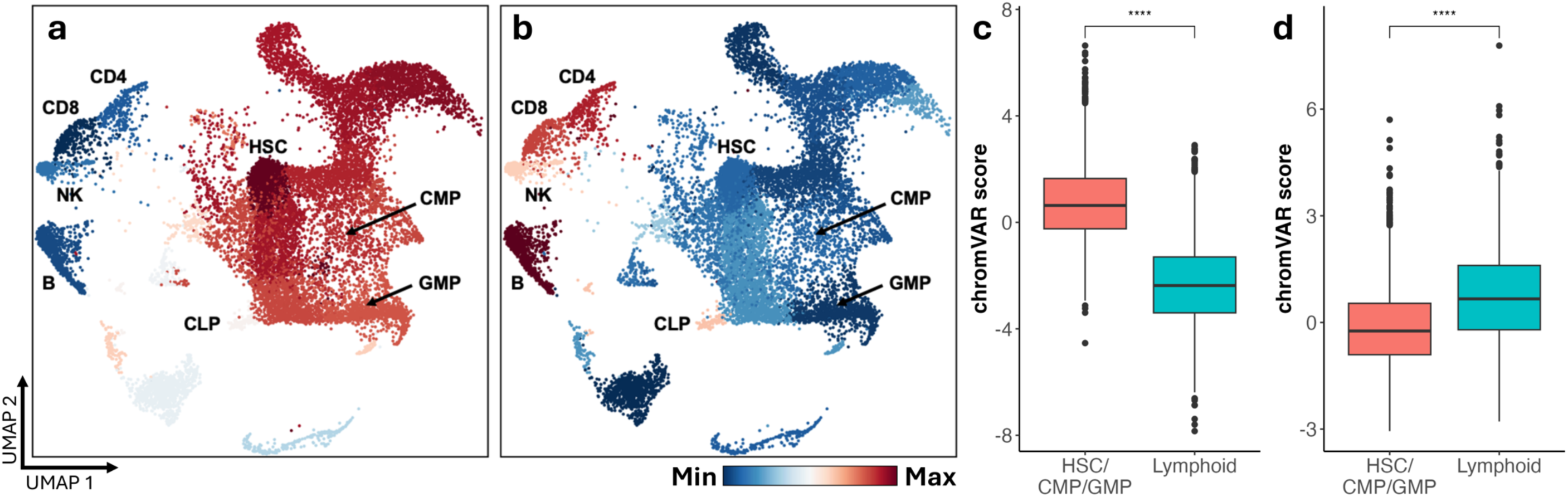
Chromatin accessibility patterns in non-malignant cells associated with lineage-enriched ESLs. **a, b,** Chromatin accessibility UMAPs of 6,088 hematopoietic cells from myelofibrosis patients (Izzo et al., 2024). Only cells that were wild type for the JAK2^V617F^ mutation were included in the analysis. Each cell is colored according to the average chromVAR deviation score for its respective cell type, based on chromatin accessibility at genomic regions corresponding to (**a**) lymphoid-enriched ESLs and (**b**) myeloid-enriched ESLs. Red shading indicates less compacted chromatin, while blue shading indicates more compacted chromatin. **c, d,** Comparison of hematopoietic stem cells (HSCs), common myeloid progenitors (CMPs), and granulocyte-monocyte progenitors (GMPs) to lymphoid cell types (CLP, B, CD4 T, CD8 T, NK) based on chromVAR deviation scores corresponding to (**c**) lymphoid-enriched ESLs and (**d**) myeloid-enriched ESLs. **** p < 0.0001 for Wilcoxon test.

**Extended Data Fig. 3:**
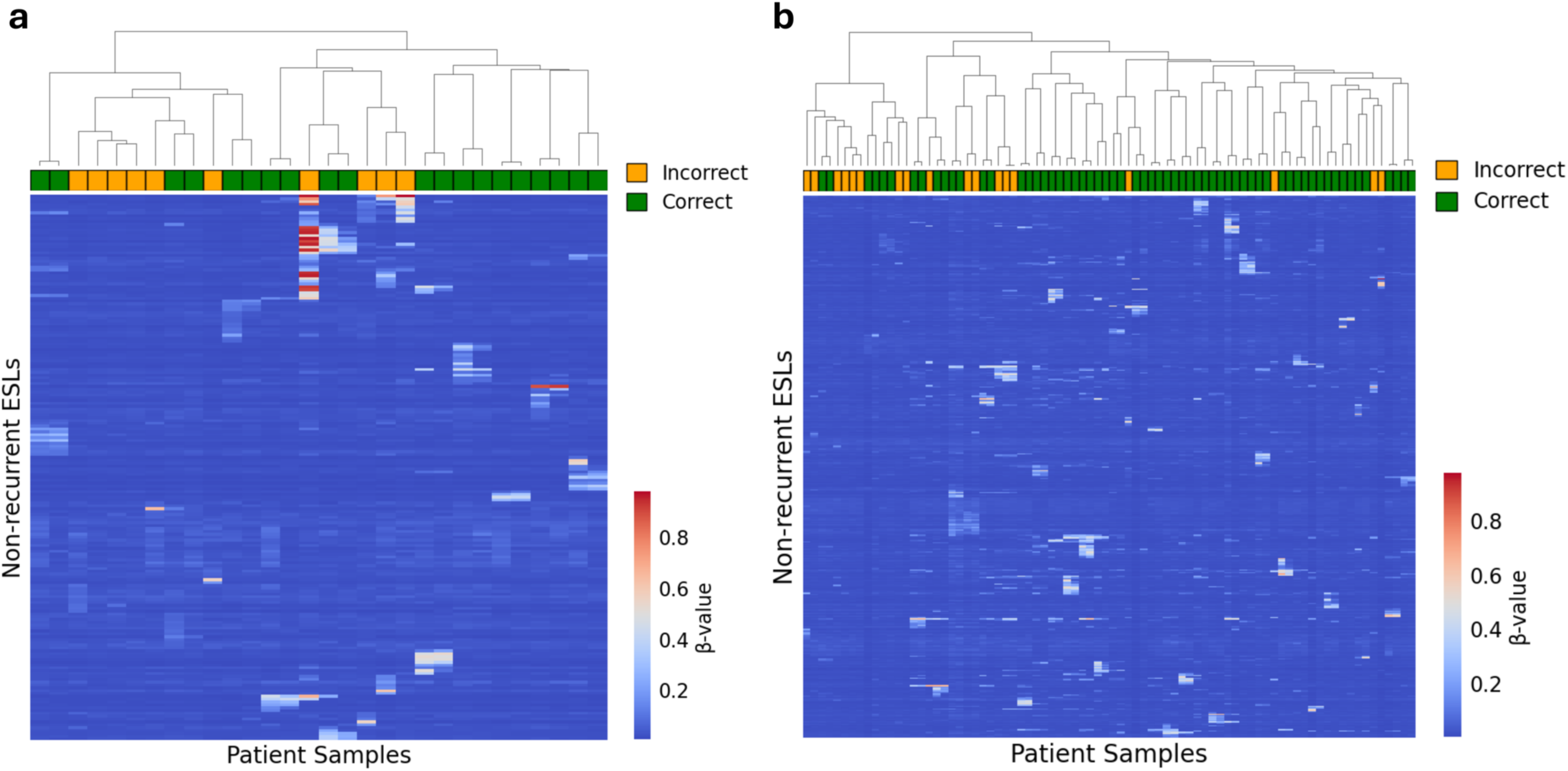
Aberrant DNA methylation in low-recurrence destabilized ESLs links diagnosis and relapse leukemic epigenetic clones. Hierarchical clustering of paired diagnosis and relapse samples from **a**, AML patients and **b**, CLL patients. The top 15 most destabilized ESLs with recurrence below 5% were selected from each patient’s relapse sample and used for clustering. Correctly paired diagnosis and relapse samples are indicated in green and incorrectly paired samples are indicated in orange. 10/15 AML patients and 31/40 CLL patients had correctly paired diagnosis-relapse samples.

**Extended Data Fig. 4:**
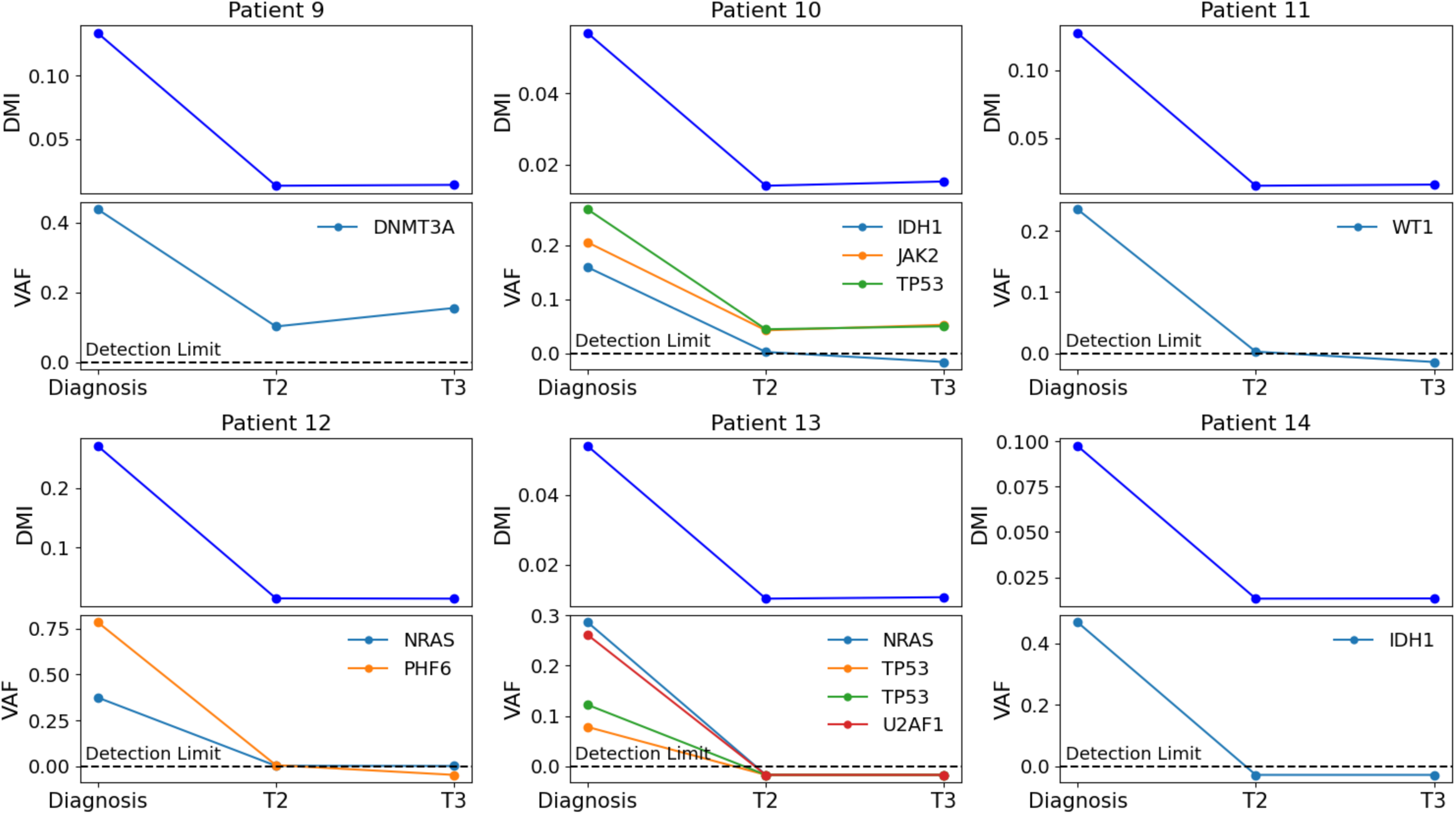
Longitudinal analysis of peripheral blood samples from 6 additional AML patients. Samples were collected at diagnosis and two subsequent time points post-therapy. The top panels display DNA methylation instability (DMI) levels, and the bottom panels show variant allele fractions (VAF) of mutations detected at diagnosis.

**Extended Data Fig. 5:**
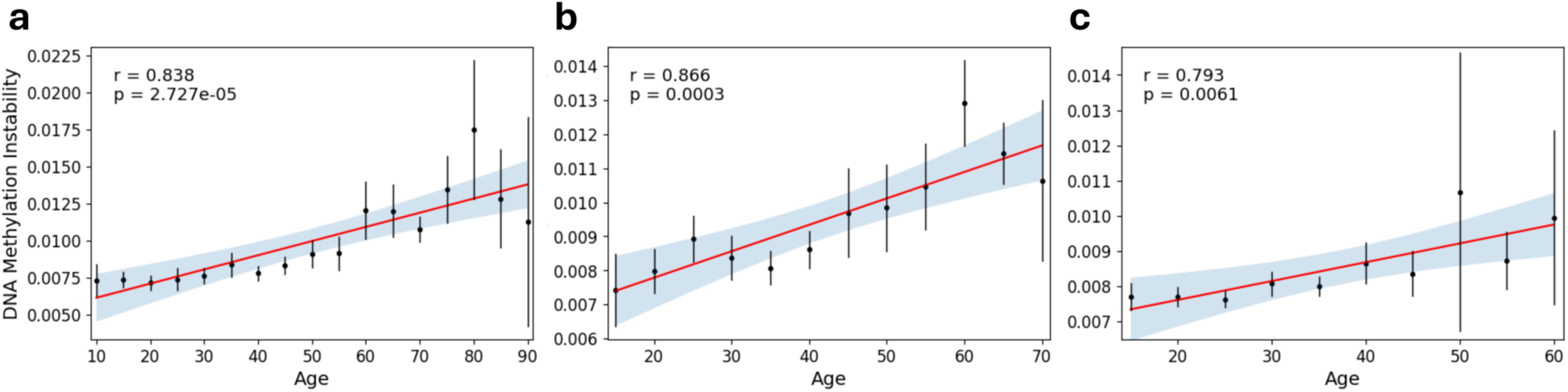
Increasing DNA Methylation instability correlates with advanced age. DNA methylation instability levels were assessed in three cohorts (n = 729, 366, 282), with participants grouped into 5-year age categories. Black dots represent the mean DMI value for each age group, with error bars indicating the 95% confidence interval. Linear regression was performed on the group means, with the regression line shown in red. The blue shading represents the 95% confidence interval of the regression line. Pearson’s correlation coefficient (r) and *P*-values are shown.

**Supplementary Fig. 1:**
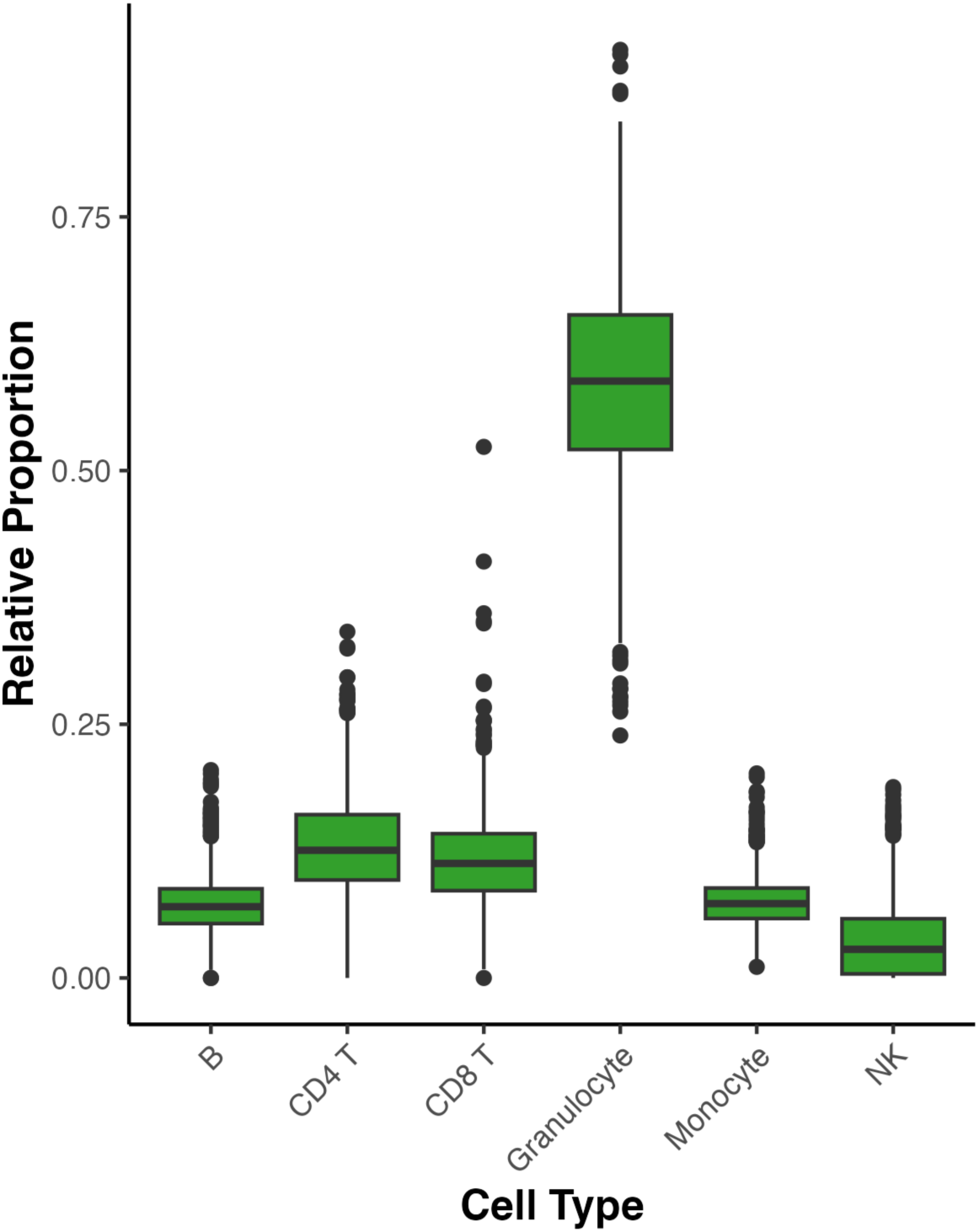
Estimation of Blood Cell Counts. Blood cell counts for all individuals in the GSE105018 cohort (n=1658) were estimated using the estimateCellCounts function from the minfi package (Aryee, M. J. et al, 2014) which infers the relative proportions of different blood cell types based on DNA methylation array data. The y-axis represents the estimated relative proportions of each blood cell type. Individuals identified as outliers — defined as values that fall outside 1.5 times the interquartile range from the first or third quartile — are shown as dots. These outliers were excluded from subsequent analyses, resulting in a final Discovery Cohort size of 1525 individuals.

**Supplementary Fig. 2:**
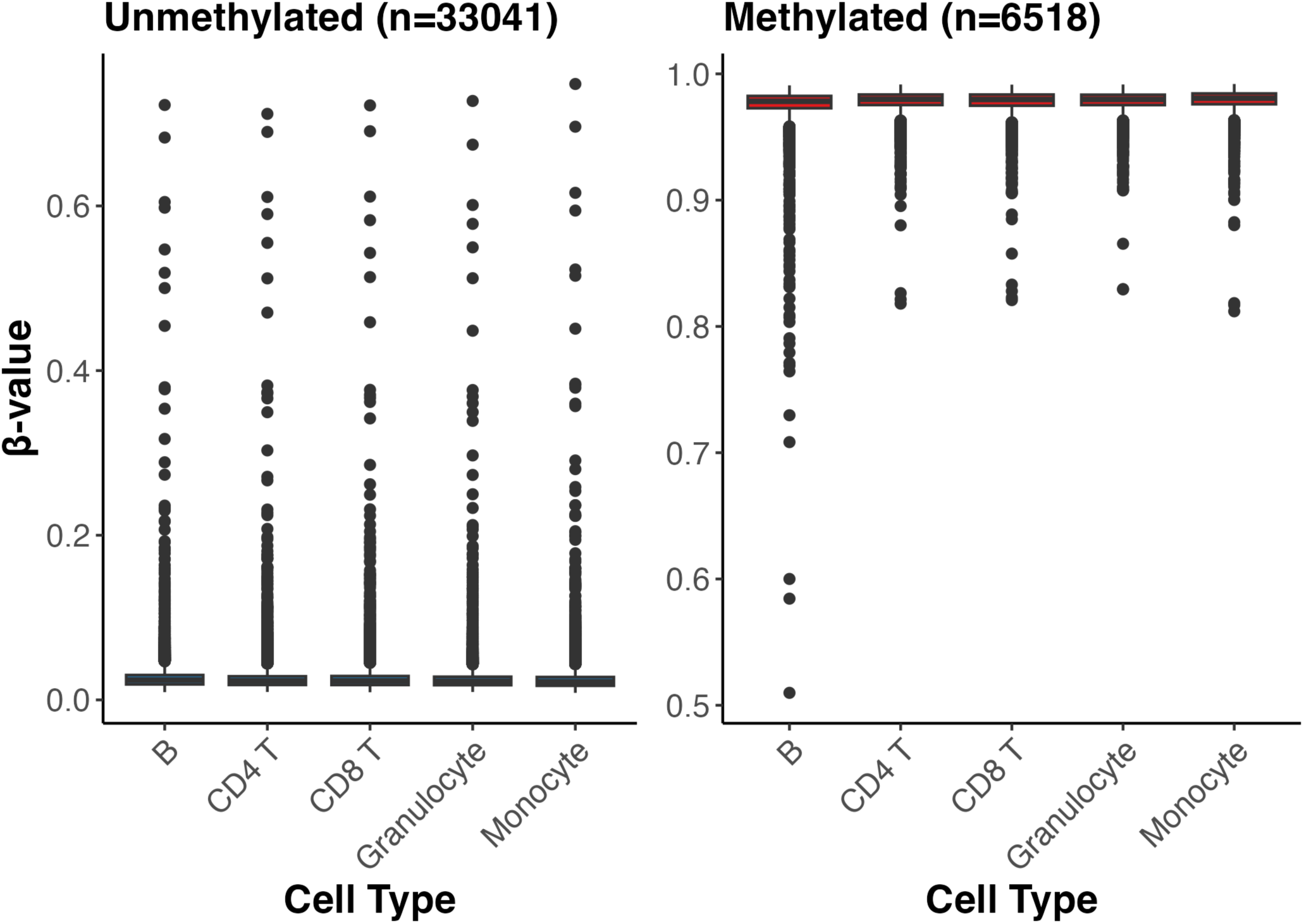
Methylation levels of highly stable sites in purified blood cell populations. Average β-values of the 10% lowest variance CpG sites from the discovery cohort were examined across five purified blood cell populations. CpGs exhibiting outlier β-values — defined as values that fall outside 1.5 times the interquartile range from the first or third quartile — are shown as dots. These outliers were excluded from subsequent analyses, resulting in a final set of 31,569 unmethylated ESLs and 6,099 methylated ESLs.

**Supplementary Fig. 3:**
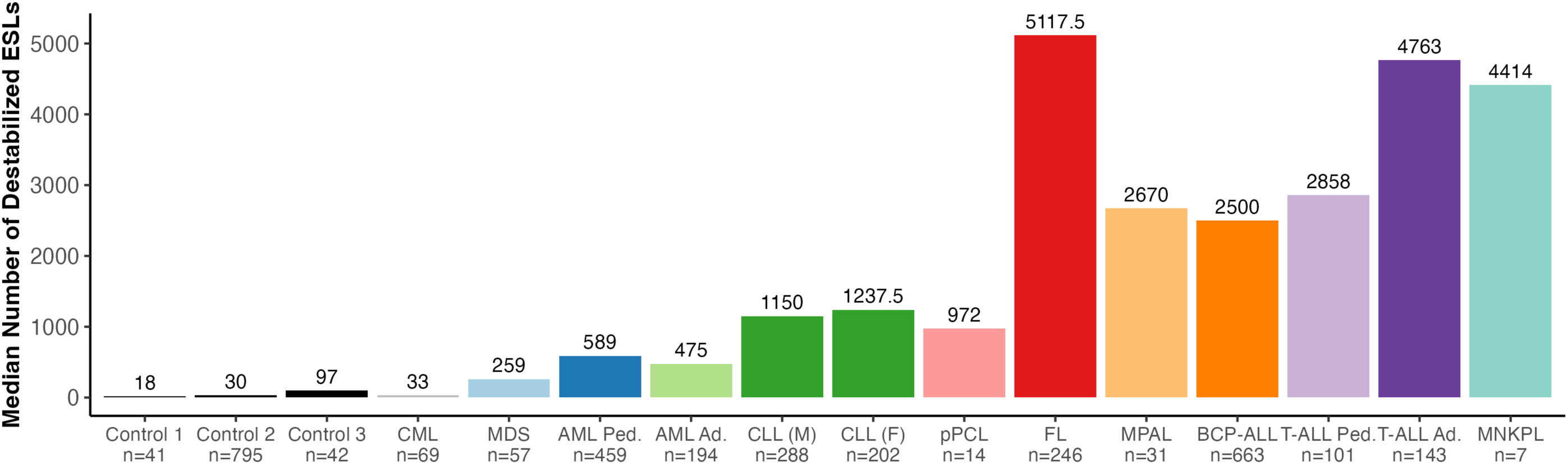
Number of Destabilized ESLs in Control and Hematological Malignancy Cohorts. For each cohort, the median number of destabilized unmethylated ESLs is shown. An ESL was classified as destabilized if its β-value exceeded the 99.9th percentile of β-values observed in the control cohorts.

**Supplementary Fig. 4:**
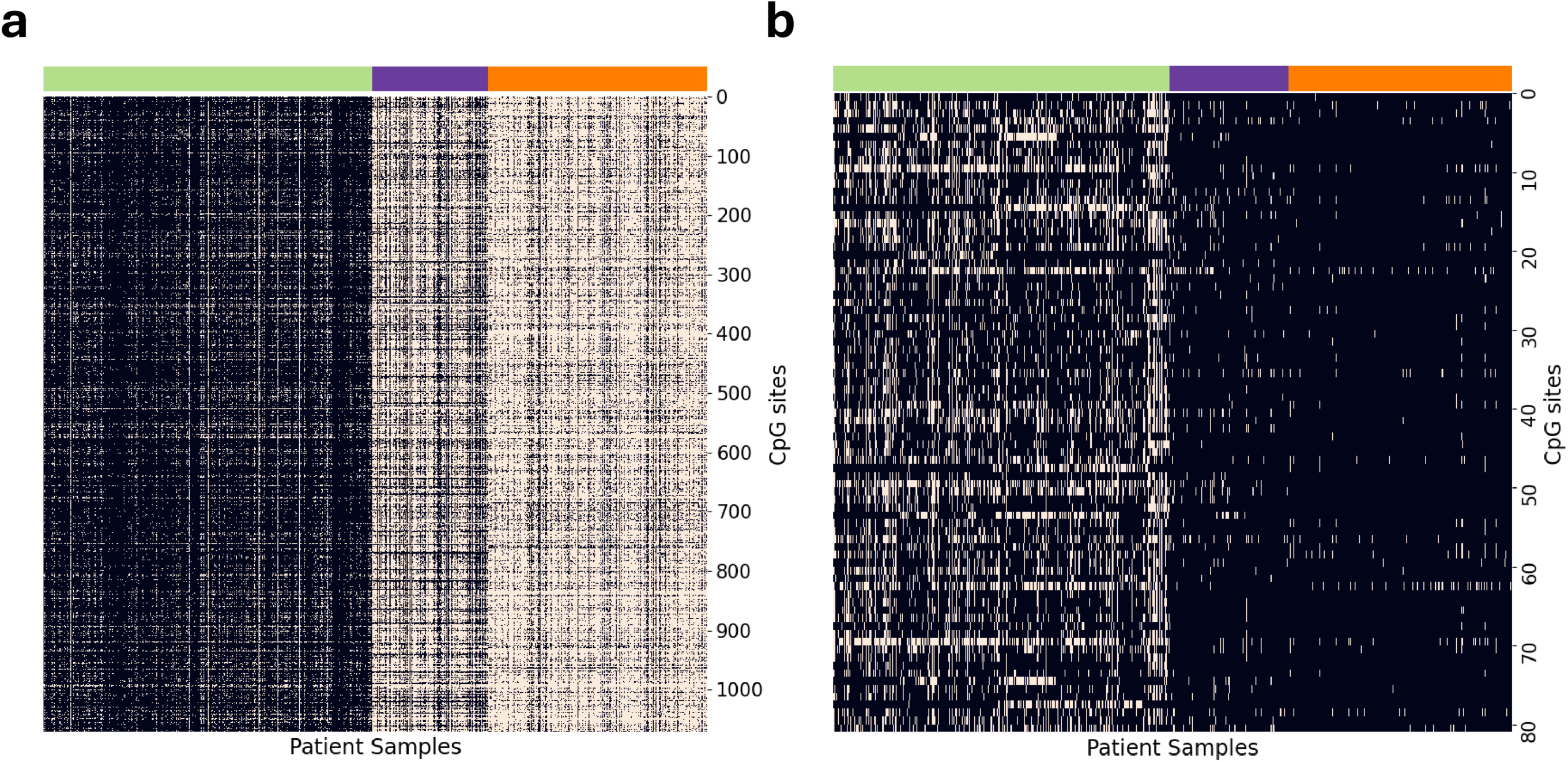
Lymphoid-enriched and myeloid-enriched ESLs. **a,** Heatmap of binarized β-values for lymphoid-enriched ESLs (n = 1072) in AML (green, n = 997), T-ALL (purple, n = 353), and BCP-ALL (orange, n = 663) patient cohorts. **b,** Heatmap of binarized β-values for myeloid-enriched ESLs (n = 81). Binarized β-values are categorized as either destabilized or stable based on the 99.9th percentile threshold derived from healthy cohorts.

**Supplementary Fig. 5:**
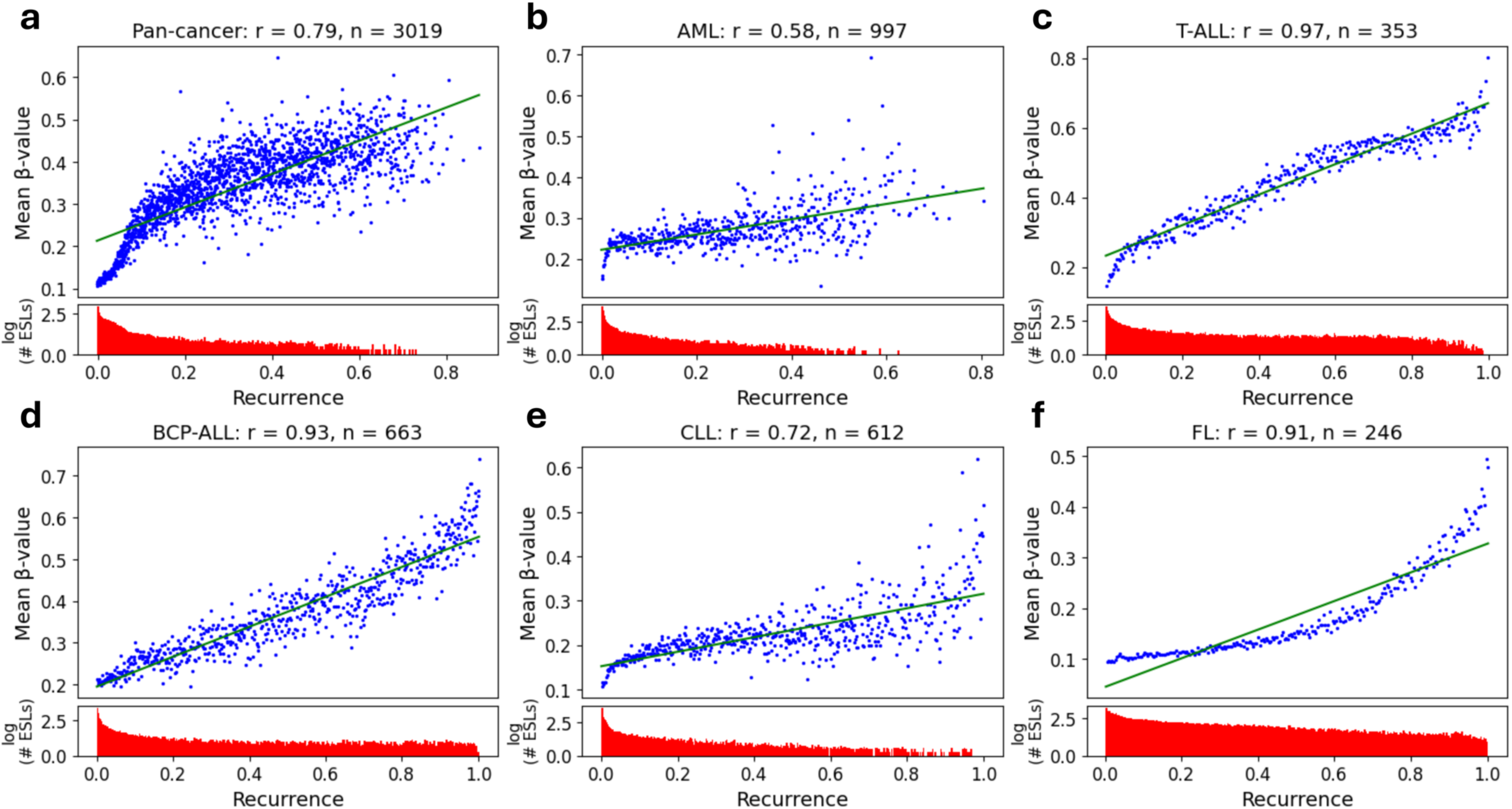
Correlation between recurrence and β-value in cancer cohorts. For each cohort, the recurrence of an epigenetically stable locus was defined as the proportion of patients in which that locus was destabilized (i.e., exceeding the 99.9th percentile threshold established from control cohorts). The y-axis corresponds to the mean β-value among patients with a given locus destabilized. Pearson’s correlation coefficient (r) and the number of samples in each cohort are shown. *P* value < 0.001 in all analyses. The bottom panels display the number of ESLs (log10 scale) across different recurrence levels. **a,** Pan-cancer analysis. **b,** AML: Acute Myeloid Leukemia, **c,** T-ALL: T-cell Acute Lymphoblastic Leukemia, **d,** BCP-ALL: B-cell Precursor Acute Lymphoblastic Leukemia, **e,** CLL: Chronic Lymphocytic Leukemia, **f,** FL: Follicular Lymphoma.

**Supplementary Fig. 6:**
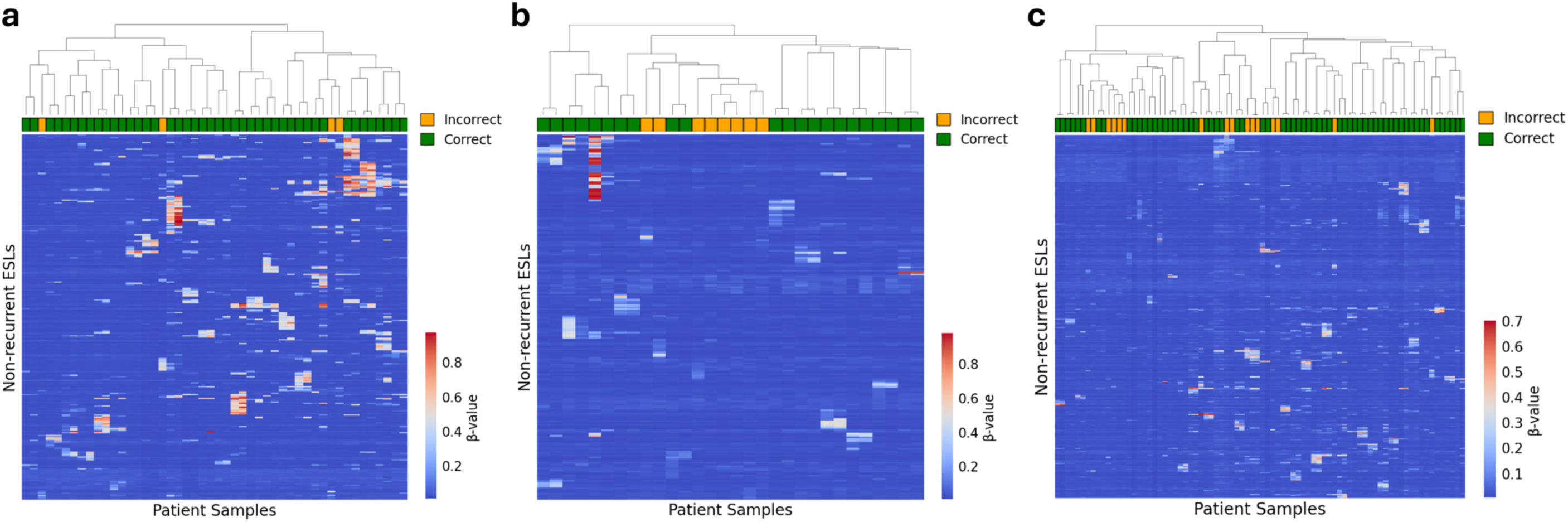
Pairing diagnosis and relapse samples using low-recurrence destabilized ESLs from diagnosis. Hierarchical clustering of paired diagnosis and relapse samples from **a**, BCP-ALL patients (n = 24), **b**, AML patients (n = 15), and **c**, CLL patients (n = 40). The top 15 most destabilized ESLs with a recurrence rate below 5% were selected from each patient’s diagnosis sample and used collectively for clustering. Correct diagnosis-relapse pairings are indicated in green and incorrectly paired samples are indicated in orange. Correct pairings were observed in 22/24 BCP-ALL patients, 11/15 AML patients, and 32/40 CLL patients.

**Supplementary Fig. 7:**
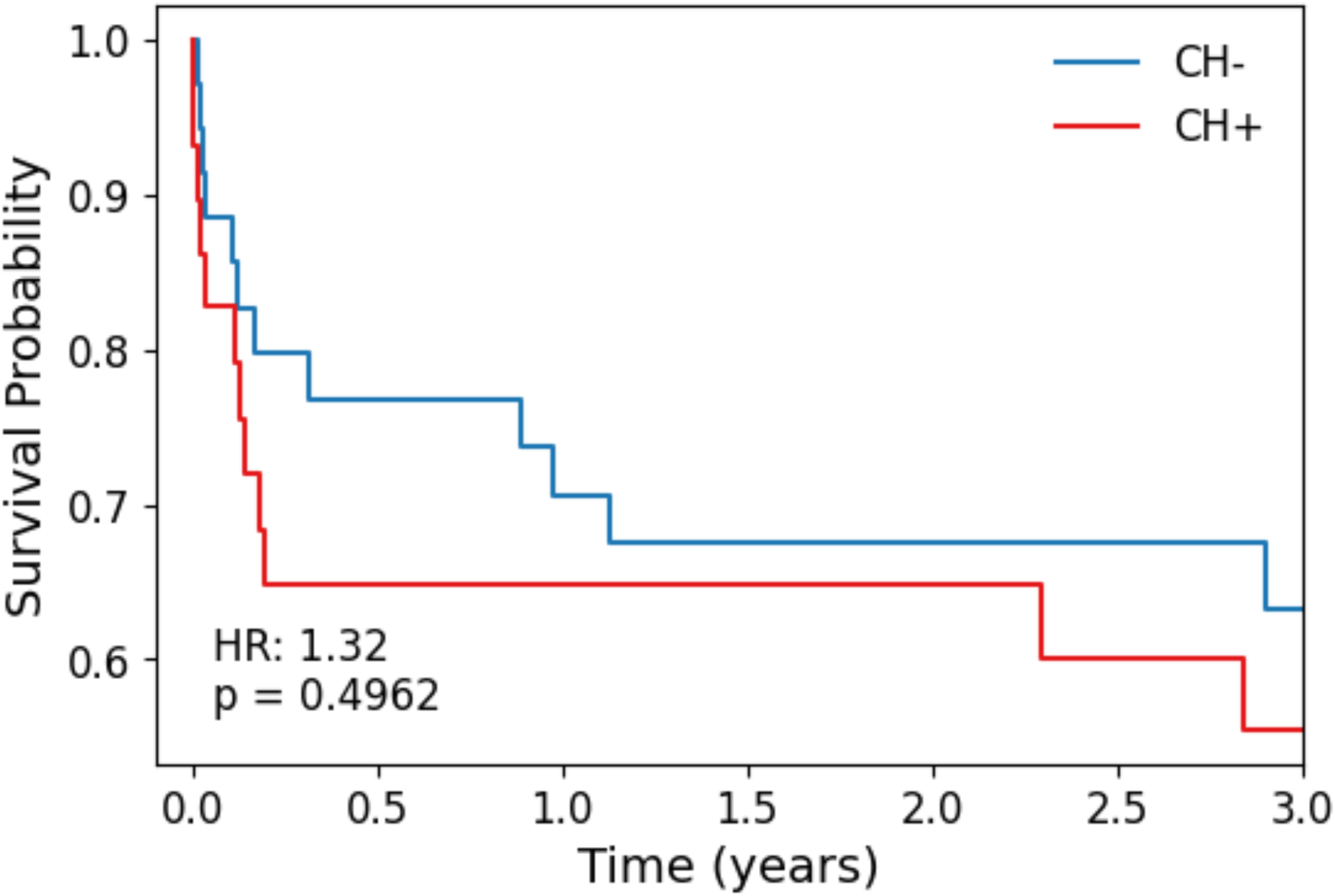
Survival analysis of the cardiogenic shock sub-cohort, stratified by clonal hematopoiesis status. Univariate Kaplan-Meier survival analysis of 64 cardiogenic shock patients, categorized by the presence or absence of clonal hematopoiesis (CH) mutations.

**Supplementary Fig. 8:**
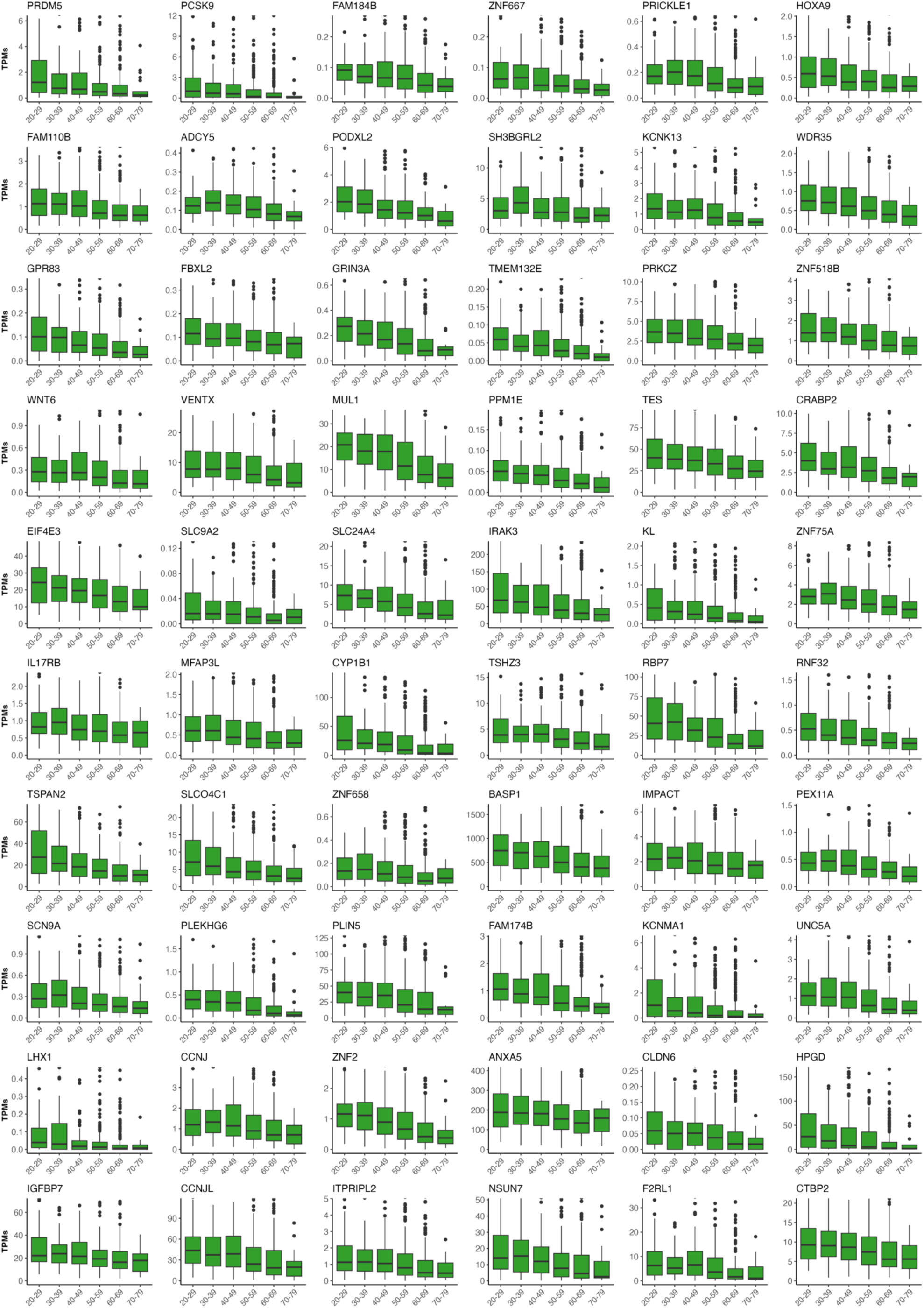
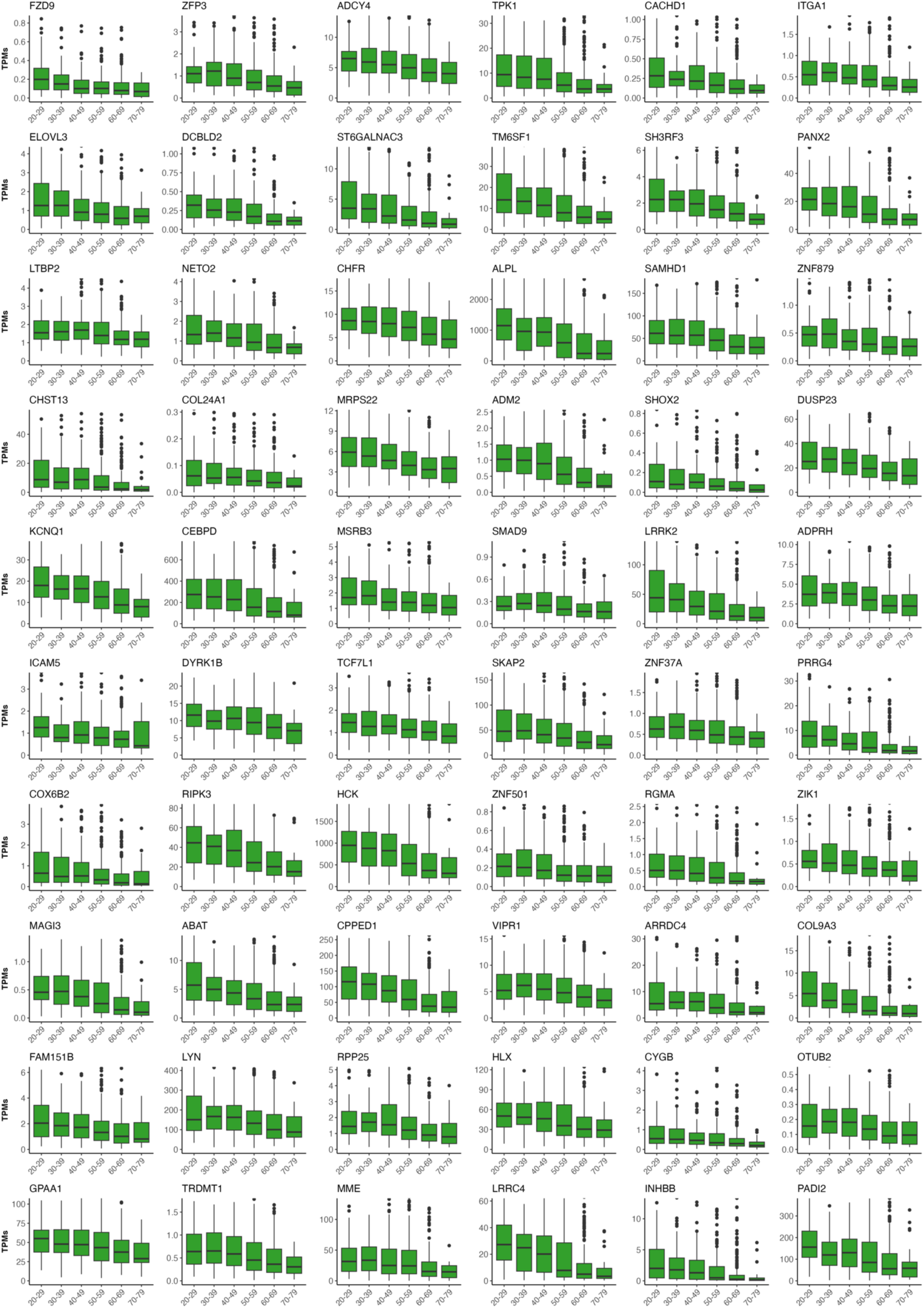
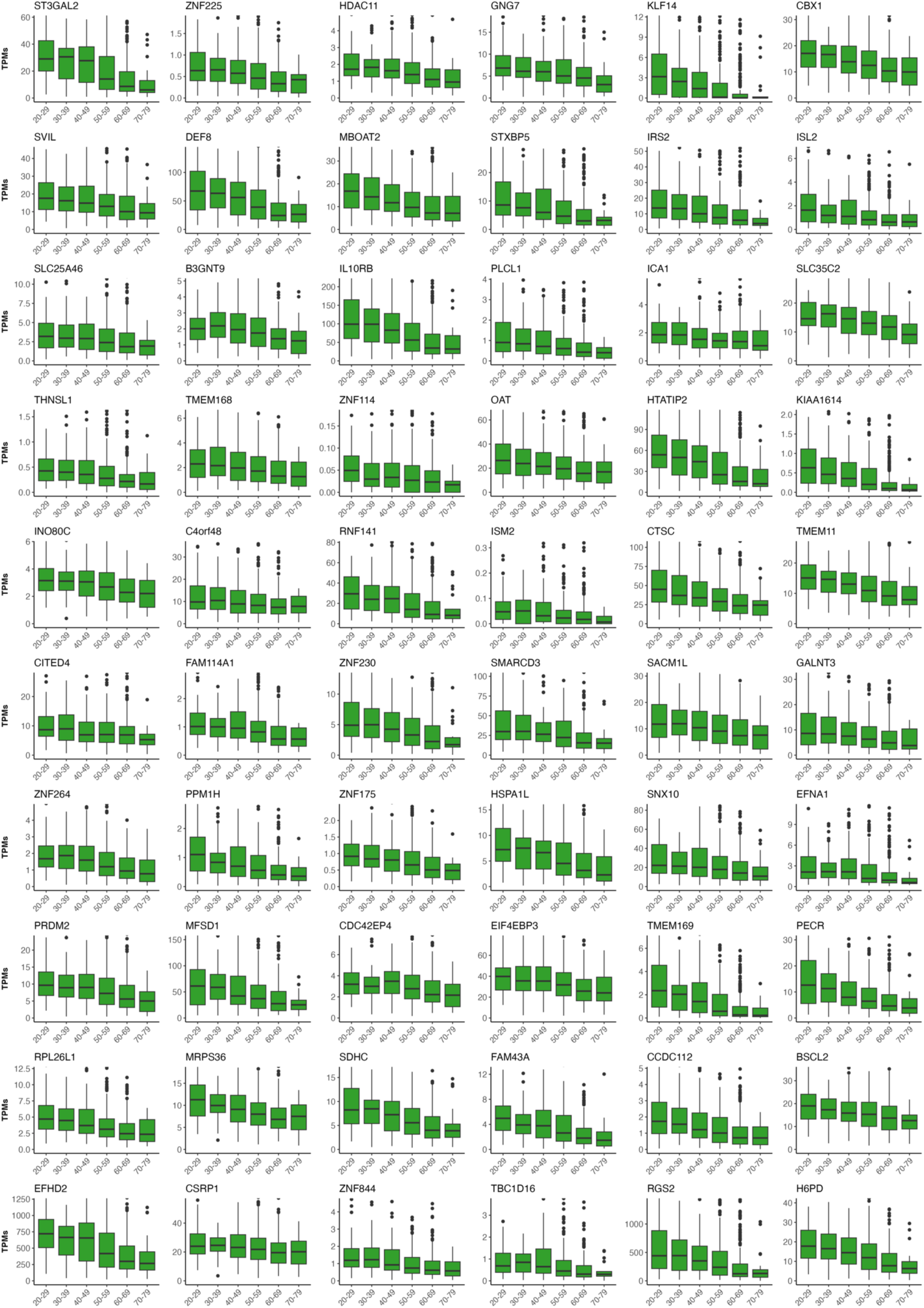
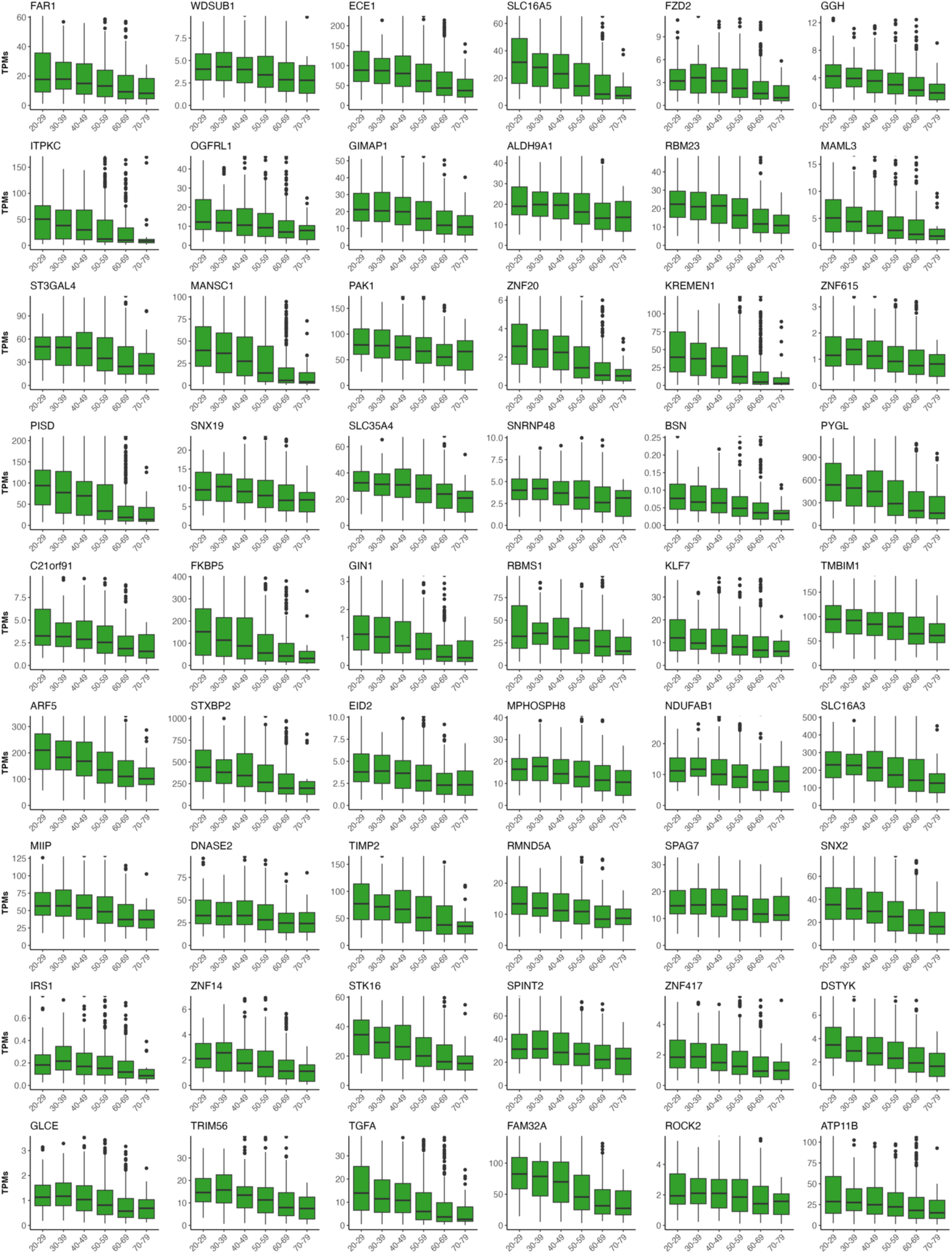
Age-dependent expression of 234 genes in peripheral blood from healthy donors. Individuals were assigned to 10-year age bins based on available data from the Genotype-Tissue Expression Project (n = 755). Y-axis represents transcript per million (TPMs). X-axis corresponds to the following age groups: 20-29, 30-39, 40-49, 50-59, 60-69, 70-79.

