## Supplementary Note 1 for "Blood-Based Epigenetic Instability Linked to Human Aging and Disease"

Supplementary Note 1: Statistical Analysis of Clustering Results

Given a situation where we need to match 24 diagnosis samples to their corresponding relapse samples (a total of 48 samples), we can calculate the probability of randomly making all the correct matches. The problem can be solved with combinatorics, where we are forming pairs from a set of items.

<u>Total Number of Possible Pairings</u>
The total number of possible pairings can be computed using the formula:
Total Pairings = (2n)! / (2^n * n!)
Where n is the number of pairs. For our case, n = 24.

Thus, the total number of ways to form pairs from 48 samples is given by:
Total Pairings = (48)! / (2^24 * 24!)

<u>Correct Pairing</u>
There is only one correct pairing, where each diagnosis sample is matched with its corresponding relapse sample. The probability of randomly matching all samples correctly is given by the ratio of the correct pairing to the total number of possible pairings:

Probability = 1 / [(48)! / (2^24 * 24!)]
This simplifies to:
Probability = (2^24 * 24!) / (48!) = 8.385265e-31

The probability of correctly matching all 24 diagnosis samples to their corresponding relapse samples by random pairing is extremely close to zero.
